# Quantitative, Multispecies Monitoring at a Continental Scale

**DOI:** 10.1101/2025.10.30.685180

**Authors:** Gledis Guri, Owen R. Liu, Ryan P. Kelly, Megan R. Shaffer, Kim M. Parsons, Ana Ramón-Laca, Krista M. Nichols, Pedro F. P. Brandão-Dias, Abigail Wells, Andrew Olaf Shelton

## Abstract

Molecular data from environmental samples can reflect the abundance of species’ DNA, an index immediately relevant to natural-resources management at a broad geographic scale. These data commonly derive from assays designed and targeted for individual species (e.g., using qPCR or ddPCR), or from metabarcoding that use more general PCR primers to amplify many species simultaneously. Multispecies analyses make efficient use of field samples and laboratory time, and speak to inherently multispecies questions of management and ecological interest. However, unlike single-species techniques, metabarcoding alone generally reflects only the environmental DNA (eDNA) proportions of target-species DNA present, not their absolute quantities. Here, we combine qPCR and metabarcoding data derived from the same samples to estimate quantities of eDNA from many fish species along the US West Coast in three dimensions, demonstrating a technique of practical relevance for both management and ecology. We derive spatially explicit maps of eDNA abundances for 12 common species of ecological and management importance and point the way to quantitative surveys of wild species using molecules alone. We find that species distribution maps derived from eDNA largely mirror known species niches and known spatial distributions. Notably, our analyses indentify biodiversity hotspots that align with previously documented regions of ecological significance, such as the Columbia River plume and the Heceta Bank.

## Introduction

Environmental DNA (eDNA, residual genetic material sampled from water, soil, or air) is an increasingly common tool for non-invasively sampling marine and aquatic ecological communities, valuable for estimating species occurrence (Veilleux et al., 2021), biodiversity (i.e., richness; Muenzel et al., 2024), and genetic structure (Andres et al., 2023). However, despite its widespread use for detecting taxa and comparing community compositions, eDNA has not been widely used for quantitative, the data upon which many practical management and conservation applications rely (but see Shelton et al. (2022), Guri et al. (2024a), and Stoeckle et al. (2022)). The use of eDNA for quantification is supported both conceptually – where there is more of a species, more cells are shed in the environment, and thus there is more DNA – and empirically where studies have reported strong linkages between single-species eDNA concentrations and organismal abundance in aquaria (Jo et al., 2019; Ledger et al., 2024), rivers and streams (Pont, 2024), estuaries and nearshore habitats (DiBattista et al., 2022; Baetscher et al., 2023), and coastal oceans (Maes et al., 2023; Shelton et al., 2019). However, uses of quantitative eDNA observations in management applications have significantly lagged behind their use of these observations for occurrence and biodiversity applications.

Surveys of marine environments play a fundamental role in supporting fisheries and conservation of protected species but are often difficult and expensive endeavors. While trawling, acoustics, or other traditional sampling methods have long been used to directly measure species abundance and biomass, they can be impractical for species inhabiting inaccessible habitats, occurring at low densities, or species that are easily injured during capture. By contrast, eDNA sampling captures information for a broader range of taxa, including those that are difficult or impossible to study with traditional gear (Ledger et al., 2025). As such, eDNA can meaningfully complement traditional survey methods for commercially important species while at the same time revealing the abundance and distribution of elusive or understudied species (Johnson et al., 2025). Notably, in 2025 the U.S.-Canada joint Pacific hake *(Merluccius productus)* stock assessment, for the first time, incorporated an eDNA-derived abundance index – an important proof of concept for integrating molecular data into formal fisheries assessments.

In recent years, substantial progress has been made toward developing eDNA as a quantitative tool for natural-resources management. Multiple studies have used species-specific assays such as qPCR or ddPCR to link eDNA concentrations (molecular abundance) with organismal abundance across commercially and ecologically important taxa, including Pacific hake, Atlantic cod, and several coastal flatfishes (Salter et al., 2019; Shelton et al., 2022; Stoeckle et al., 2022; Guri et al., 2024a). Although these studies have concluded direct correlation between molecular and biomass-based abundances, it is important to note that such relationships tend to hold at regional scales rather than at individual sampling stations. This pattern likely reflects the influence of biophysical transport and mixing, as eDNA is advected and dispersed over distances of only a few kilometers before degradation and dilution attenuate the signal (Collins et al., 2018; Guri et al., 2024a; Xiong et al., 2025). However, single-species approaches remain impractical for large-scale applications because each species requires independent assay development, optimization, and processing (Ruppert et al., 2022; Langlois et al., 2021).

In contrast, multispecies analysis (metabarcoding) reflects information on many species of interest simultaneously, but at the costs of producing compositional estimates of proportional abundance (Gloor et al., 2017; McLaren et al., 2019) that may require statistical correction for species-specific biases due to PCR amplification (Shelton et al., 2023). Consequently, metabarcoding data alone – without calibration to independent quantitative measures such as qPCR, mock communities for alleviating PCR biases, or independent traditional survey data – cannot yield information about the abundance of DNA from environmental samples (e.g. Guri et al. (2024a)). Nonetheless, metabarcoding’s ability to capture information across diverse taxa offers a promising path toward ecosystem-level monitoring essential for modern fisheries and conservation management. As agencies begin to implement the new National Aquatic eDNA Strategy (Kelly et al., 2024)– the first goal of which is to fold eDNA data into federal decision-making – this and other large-scale examples point the way to maximizing the benefit of this information-rich data source for natural-resources management at national scale.

Although a few recent studies have calibrated metabarcoding results against quantitative or independent abundance measures, these efforts have so far been confined to small regional scales (Stoeckle et al., 2022; Guri et al., 2024a), highlighting the need for methods that extend such approaches to broader, management-relevant spatial domains.

Here, we develop quantitative estimates of eDNA abundance for a diverse group of fish species spanning different depth niches across 10 degrees of latitude along the US West Coast entirely from eDNA. We integrate single- and multi-species (using MiFish-U universal primers (Miya et al., 2015)) techniques to create three-dimensional distribution maps, leveraging models established in previous studies (Shelton et al., 2022; Guri et al., 2024a; Shelton et al., 2023; Allan et al., 2023). By calibrating metabarcoding data – correcting raw species observations to their initial proportions using the known concentration of a list of species from tissue samples (hereafter mock community; Shelton et al., 2023) – and using the absolute eDNA quantity of a reference species (here Pacific hake (*Merluccius productus*)), we expand initial estimated proportions into absolute estimates of the abundance of each species’ eDNA (Guri et al., 2024a). We subsequently smooth the estimated eDNA quantities to generate distribution maps (Liu et al., 2023). We present quantitative results for 12 fish species using eDNA, comparing them with literature-based species distributions to demonstrate the validity and applicability of our approach, but note that the method immediately generalizes to any ecological community of interest. This study aims to show that the eDNA-derived estimates presented here are conceptually analogous to the unitless abundance index used in fisheries stock assessments – measures proportional to biomass – and are therefore inherently compatible with existing assessment frameworks, requiring minimal modification for integration (Johnson et al., 2025).

## Methods

### Sample Collection, Processing, and Quantitative PCR (qPCR)

We used seawater samples collected in 10 L Niskin bottles deployed on a CTD rosette from dusk to dawn at five depths (0, 50, 150, 300, 500 m) from stations along the West Coast of the US aboard the ship *Bell M. Shimada* during Summer 2019 on the Joint U.S. – Canada Integrated Ecosystem and Pacific Hake Acoustic-Trawl Survey (de Blois, 2020). 2.5 L water samples were collected from each Niskin and filtered immediately on 47 mm diameter mixed cellulose-ester sterile filters with a 1 *µ*m pore size, subsequently extracted using a phase-lock phenol-chloroform protocol, and analyzed in triplicate using a qPCR assay, multiplexed for Pacific hake, sea lamprey, and eulachon, as described in Ramón-Laca et al. (2021) and Shelton et al. (2022). A subset of the samples (n = 554) collected in Shelton et al. (2022), was used for metabarcoding analysis in this study in addition to qPCR samples already published for hake quantification in Shelton et al. (2022) (n= 1,794).

### Metabarcoding: Environmental Samples and Mock Community

In total, 554 environmental samples were randomized and sequenced across 7 Illumina MiSeq (v3 600 cycle kit) libraries. Total PCR reactions, including PCR blanks (n = 7) and positive controls (n = 7), were amplified using MiFish-U universal primers (Miya et al., 2015) (see SI Appendix Extended Methods for details on metabarcoding methods). Importantly, we metabarcoded only one water sample collected from each sampling station.

Sequences were de-multiplexed and the adapters removed by Illumina processing software tools. Primers were removed with Cutadapt v4.9 (Martin, 2011). We then used DADA2 (default parameters; Callahan et al., 2016) to denoise sequences, remove chimeras, and generate amplicon sequence variants (ASVs). The sequences were then BLASTed against the NCBI nucleotide (nt) database (access: August 2024) using BLASTn algorithm with a cut off at 97% identity, assigning the least common ancestor of the top hit, resulting in species, genus, family, or higher rank classifications. Samples with fewer than 1000 total reads were filtered as an indicator of poor reliability. Additionally, samples lacking a positive detection of Pacific hake (reference species) in both qPCR and metabarcoding datasets were removed from downstream analysis, due to the model dependance of the reference species being present in both (see SI Appendix Extended Methods). Out of the initial environmental samples, 535 passed the quality filtering and 371 of those contained Pacific hake in both data streams (see SI Appendix, Fig. S3). Details on bioinformatic pipeline are available in SI Appendix Extended Methods.

To calibrate metabarcoding observations and account for species-specific amplification bias (Shelton et al., 2023; Gold et al., 2023), we constructed multiple mock communities comprising of genomic DNA from a total of 39 fish species. Vouchered DNA extracts or tissues were obtained from either the University of Washington Fish Collection at the Burke Museum or the NOAA Northwest Fisheries Science Center collections. After quantifying the concentration of mitochondrial DNA template in each DNA extract, we constructed a total of eight mock communities of varying species compositions (see SI Appendix, Table S1). We selected 12 species (additional to Pacific hake) of commercial and ecological importance that co-occurred in the environmental samples and in the mock community for downstream analysis: Pacific herring (*Clupea pallasii*), northern anchovy (*Engraulis mordax*), northern smoothtongue (*Leuroglossus stilbius*), Dover sole (*Micros-tomus pacificus*), Pacific sardine (*Sardinops sagax*), Pacific chub mackerel (*Scomber japonicus*), widow rock-fish (*Sebastes entomelas*), northern lampfish (*Stenobrachius leucopsarus*), longfin dragonfish (*Tactostoma macropus*), plainfin midshipman (*Tarletonbeania crenularis*), eulachon (*Thaleichthys pacificus*), Pacific jack mackerel (*Trachurus symmetricus*). After amplification and sequencing (SI Appendix Extended Method), we used the mock communities (which were run in the same PCR conditions as the environmental samples) to estimate species-specific amplification efficiencies and correct for amplification bias following (Shelton et al., 2023).

### Statistical analysis

To generate estimates of DNA concentration, we constructed a joint statistical model for qPCR and Mifish-U metabarcoding observations from the same samples (Fig. 1). We used qPCR observations of hake DNA concentration for all water samples available (1,794 unique water samples were assayed and using a total of 5,394 qPCR reactions). This model takes observations of the PCR cycle at which amplification was detected in each field sample and estimates a hake DNA concentration for each water bottle collected at each sampling station-depth combination (Fig. 1). This statistical model accounts for sample-specific effects (e.g., dilution of the sample before PCR amplification) and the effect of replicate water samples (a random effect), to provide estimates of hake DNA concentration in from each water sample collected at each location and depth (Fig. 1). This qPCR component of the joint model provides a quantification of DNA concentration for one species (Pacific hake) that can be connected to the multispecies proportions provided by DNA metabarcoding (Fig. 1).

**Figure 1:**
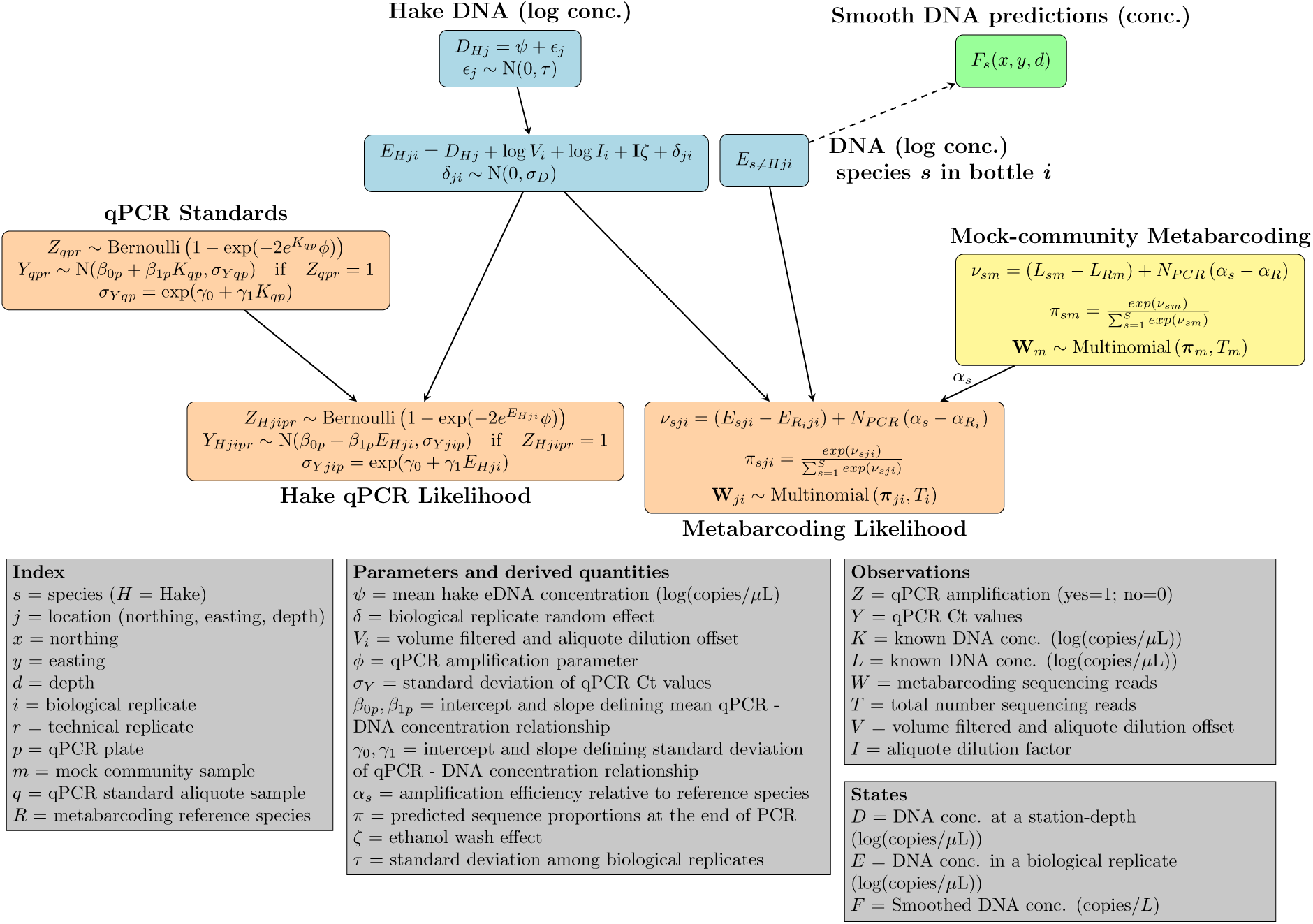
Schematic for the statistical models to produce species distributions. Four sets of observations (orange) were used simultaneously to provide estimates of eDNA concentration for twelve fish species (blue) from which we derive smoothed predictions of DNA concentration (green), and mock communities for metabarcoding (yellow) were estimated externally and provided estimated *α_s_* that were used in the joint model. Notation for parameters, subscripts, and observations are indicated in gray.

Unlike single-species observation methods, metabarcoding data are compositions, so each individual sample provides only information about species eDNA proportions rather than absolute DNA concentrations. Furthermore, metabarcoding does not yield a consistent, species-specific detection probability (i.e., sensitivity), because the likelihood of detecting a particular species depends upon the other template sequences present during PCR amplification (Shelton et al., 2023), as well as on the total sequencing depth for that sample (Kelly et al., 2019). Greater sequencing depth generally increases the probability of identifying rarer species, conditional on their DNA being collected and amplified with the chosen PCR primers.

We follow Shelton et al. (2023) and construct a compositional model for the metabarcoding data from 371 of the water samples (after quality filtering) analyzed for Pacific hake qPCR. We use the metabarcoding observations of the 8 mock communities to calibrate the relative amplification efficiency of the 12 focal species (plus hake, the 13*^th^* species). In the absence of qPCR data, this model would produce estimates of proportional DNA contributions for each water sample. But for each sample in which we have non-zero estimates of hake DNA concentration from qPCR, and non-zero estimate of hake proportional contribution from metabarcoding (SI Appendix, Fig. S3), we can expand from the DNA concentration of hake to provide an estimate of the DNA concentration of the 12 remaining focal species (Allan et al., 2023). Our model generates estimates of DNA concentration (copies/L) for each species. Hake was present in 86.7% of samples via qPCR, and 67.7% of samples via metabarcoding, making it an appropriate and important reference species compared to other species present in our data. To enhance sample coverage – by increasing the number of non-zero estimates of reference species – and improve model accuracy, multiple single-species qPCR assays (i.e., multiple reference species) could in the future be integrated as additional sources of DNA concentration. However, this approach must be applied thoughtfully, ensuring that the additional selected qPCR target species have minimal co-occurrence with the initial one, like Pacific sardine or northern anchovy in our case (SI Appendix, Fig. S4). Nevertheless, samples lacking hake detections here were sporadically distributed across the study area (SI Appendix, Fig. S3), and deserve further attention in future work.

We focused on species that were relatively common in the raw dataset – those well above the lower detection limits under our sampling and sequencing conditions – ensuring greater confidence in their consistent detectability. Although many more species were detected in the raw data, the majority were much rarer and thus less reliably observed.

The above statistical model (Fig. 1) provides predictions of species-specific DNA concentrations at each location and depth sampled (*d* = 0, 50, 150, 300, or 500 m). We used the sdmTMB package (Anderson et al., 2022) to generate smooth maps of DNA concentration for each species independently (Fig. 1, green box).

As Pacific hake (*Merluccius productus*) has been examined elsewhere using qPCR (Shelton et al., 2022), we did not include the spatial patterns of that species in our analysis.

Analyses, figure preparation, and data curation were performed in R (R Core Team, 2024) and the joint statistical model for qPCR and metabarcoding was coded in stan (Stan Development Team, 2023). For a simplified version of this model, the QM R package can be utilized (www.github.com/anonymous).

## Results

Our rigorous quantitative framework and use of hundreds of water samples provides ecological inferences at the scale of management. We find spatial patterns of DNA concentrations reflect the distributions of commercially important coastal-pelagic fish species in three dimensions and across ten degrees of latitude in a single year of sampling in 2019 (Fig. 2: northern anchovy, Pacific sardine; SI Appendix, Fig. S1: Pacific herring and jack mackerel). For other high-biomass species otherwise lacking well-described distributions, we provide the first species distribution models (Fig. 2, SI Appendix, Fig. S1). We find that the estimated eDNA concentrations are correlated among species with similar ecological traits and habitat preferences (Fig. 3). Emergent patterns of species richness and estimated eDNA aggregation correspond with known ecological drivers, evidence that eDNA traces can reliably capture ecologically meaningful patterns (Fig. 4). Particularly for forage fish species – which are critical high-biomass links in marine food webs, but difficult to survey with traditional methods (Pikitch et al., 2012) – molecular methods offer practical and quantitative indices of abundance over large spatial scales. Our work advances using the use of eDNA beyond previous research that has used eDNA for demonstrating qualitative multispecies patterns (e.g. Cornelis et al. (2024)) or quantitative analysis of single species at small spatial scales (e.g. Baetscher et al. (2024)), into a quantitative framework for multiple detected species.

**Figure 2:**
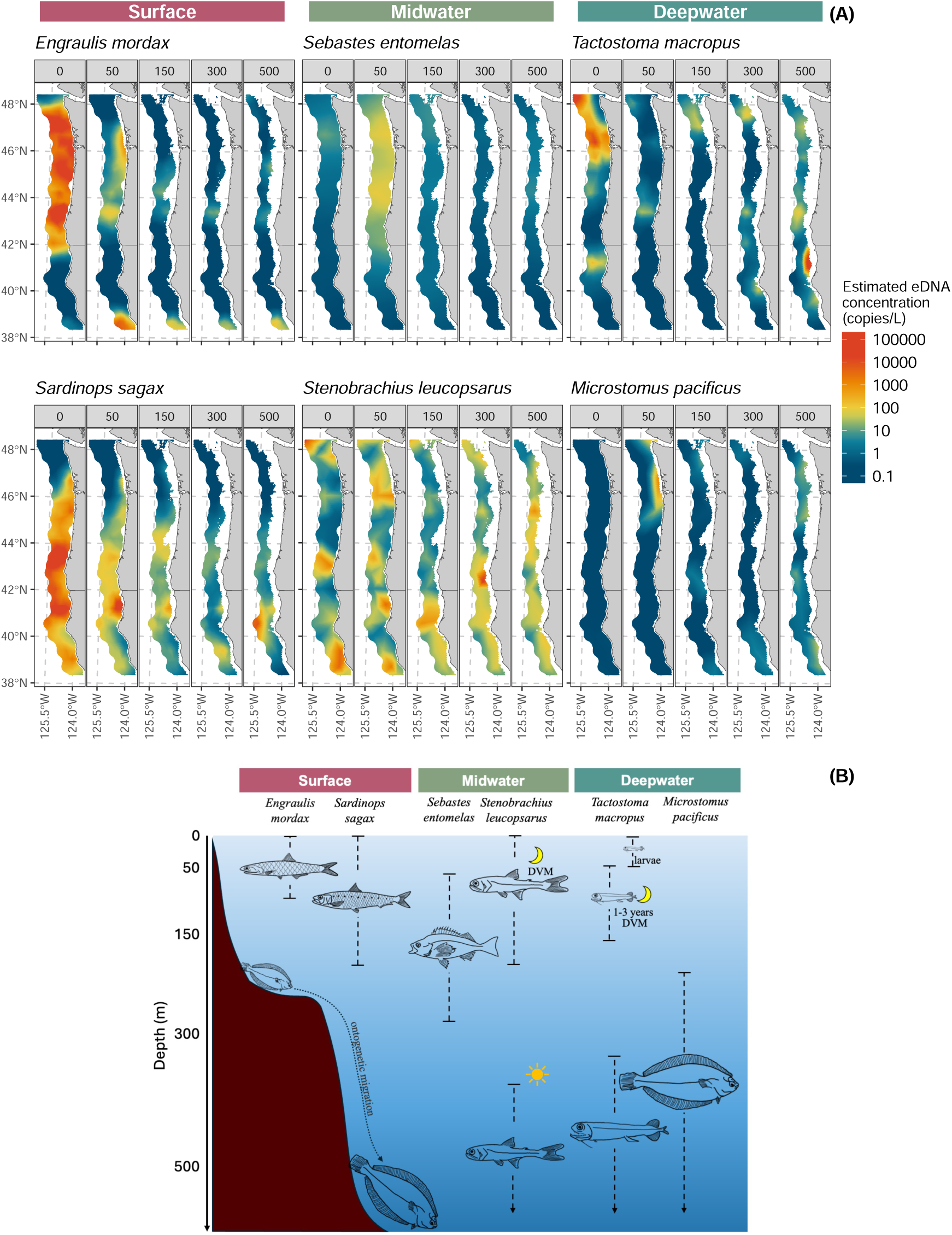
Estimated species eDNA concentrations (n=6; for the remaining species see SI Appendix, Fig. S1) from samples collected across 0, 50, 150, 300, and 500 m (A), and depth distribution of those species from the literature (B).

**Figure 3:**
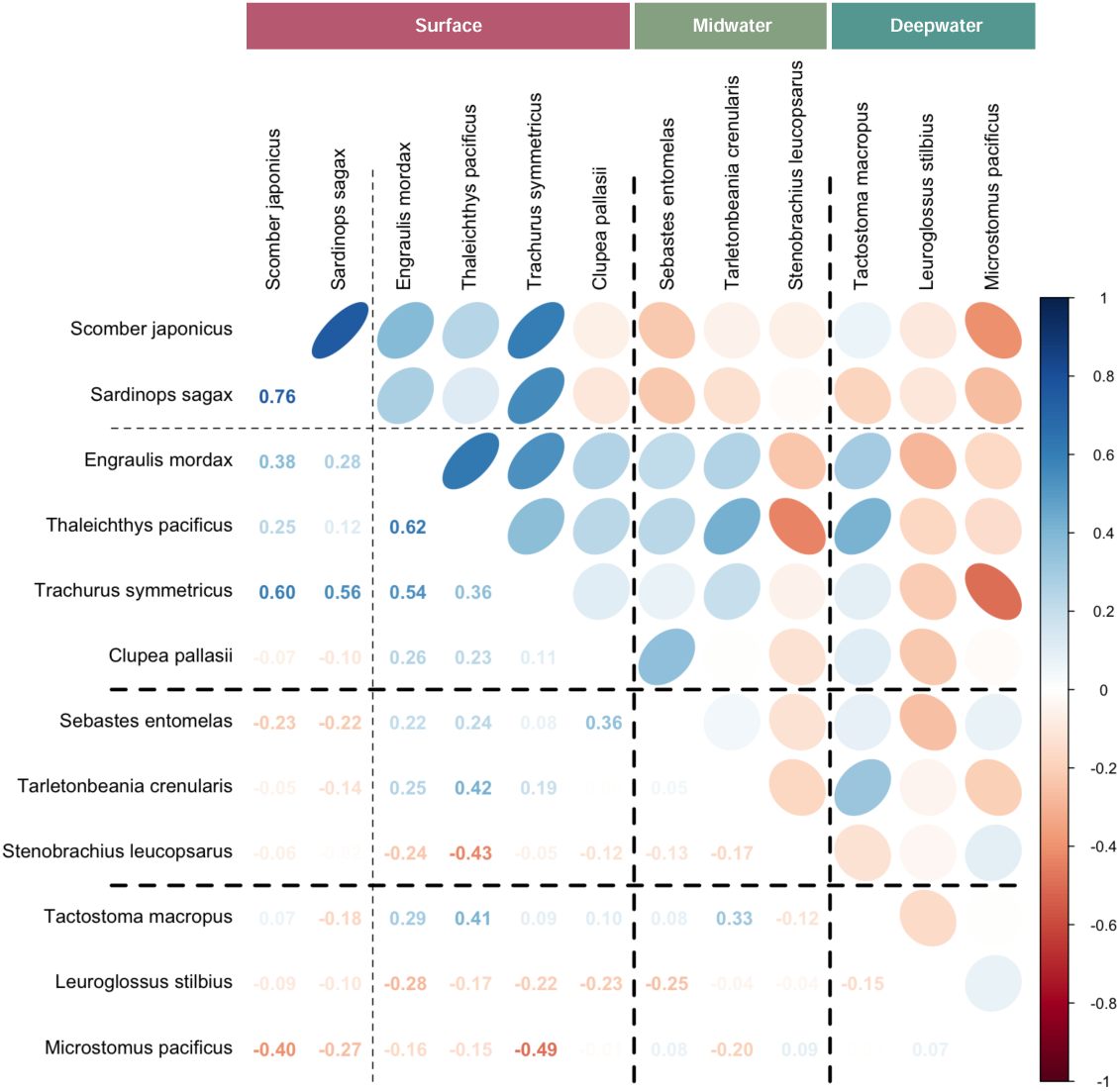
Correlation plot describing species co-abundance and co-occurance. Spearman’s *ρ* correlation coefficient are expressed in numbers and colors ranging from -1 (red) negatively correlated to 1 (blue) positively correlated. Species are ordered in known ecological groups (surface, midwater, and deepwater) to visually indicate the gradual change of species co-occurance.

**Figure 4:**
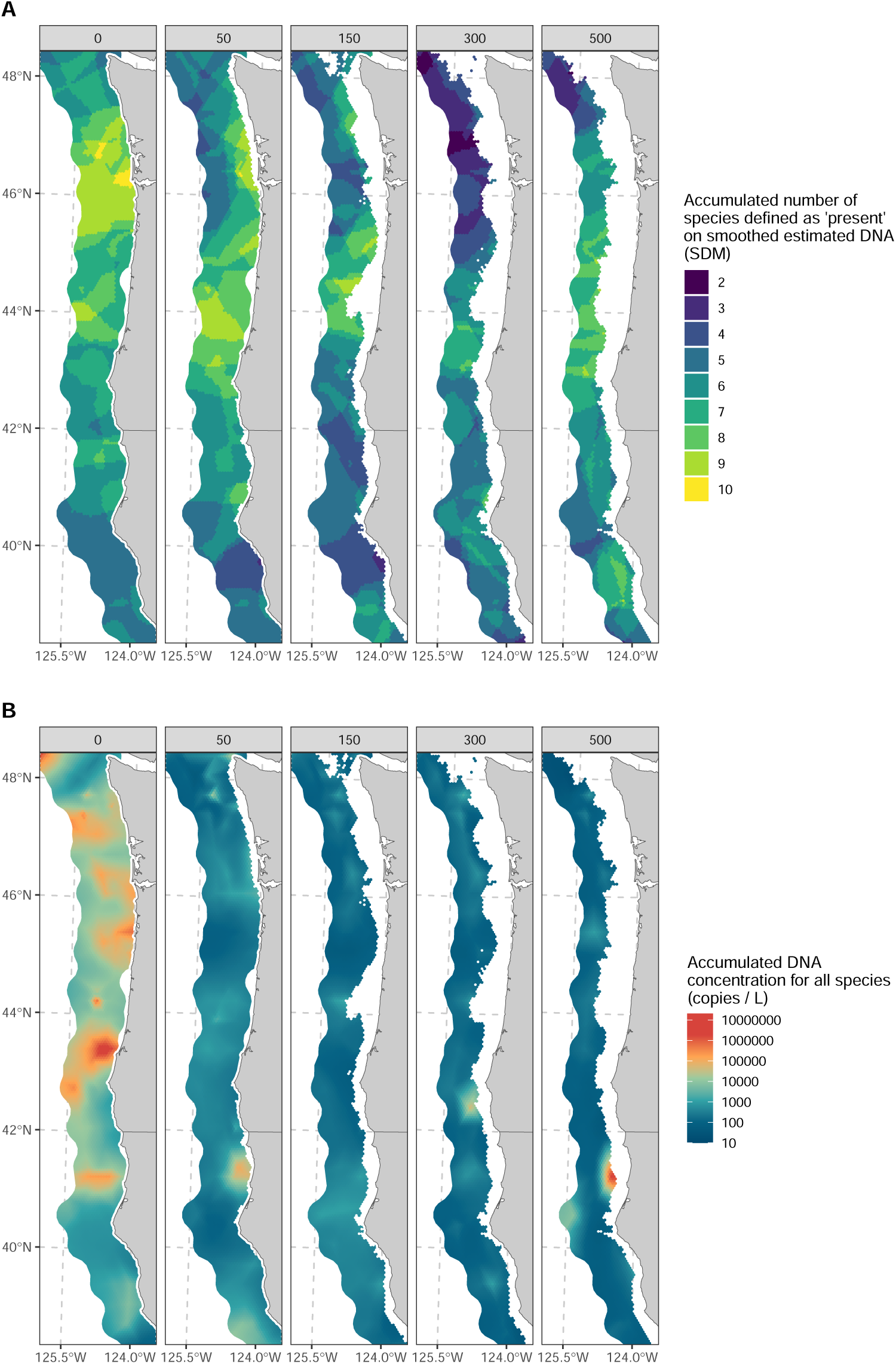
Spatial and vertical distribution of species richness (A) and environmental DNA concentration (B) across depths (0, 50, 150, 300, and 500 m).(B) Species richness represented as the cumulative number of species present (species with higher than 1.5 copies/L (see SI Appendix, Fig. S2 for defining presence) on smoothed estimated eDNA concentrations) and accumulated eDNA concentration represented as the cumulative sum of eDNA (copies/L) across all species.

Below we highlight results for six species (Fig. 2) representing a range of habitat associations and general species information; distribution maps for all the remaining species are presented in SI Appendix, Fig. S1. In particular, our analyses present detailed, depth-specific information on eDNA abundance – likely a proxy for the abundance of the fish themselves – that is difficult to obtain with traditional sampling methods.

### Surface species

Northern anchovy (*Engraulis mordax*) and Pacific sardine (*Sardinops sagax*) are two iconic coastal pelagic species of the eastern North Pacific (Steinbeck, 1945) and play a vital ecological role linking lower and upper trophic levels (e.g., diets for Chinook salmon and sea lions; Kaplan et al., 2019); historically these supported large commercial fisheries particularly in California. While larvae and juveniles typically occupy the upper 50m, adults move to 100m during the day to avoid predators (Litz et al., 2008). Our eDNA estimates consistently reflect the expected three-dimensional distributions of known surface dwelling species, with high concentrations in surface waters for both species (Litz et al., 2008; Parnel et al., 2008), and an accumulation pocket near the Columbia River outflow (46.2*^◦^*N; Fig. 2). Sardine favors slightly warmer temperatures and more southerly waters than anchovy (SST range from literature: 13–24*^◦^*C for sardine vs. 10–14*^◦^*C for anchovy). eDNA patterns tracked these differences with sardine eDNA in greatest abundance at 41*^◦^*N (range 40*^◦^*–45*^◦^*), while anchovy eDNA abundance was greatest at 43.5*^◦^*N (range 42*^◦^*–48*^◦^*; Fig. 5) such that eDNA captured subtle yet meaningful differences in habitat preferences. Additionally, eDNA abundance patterns suggested a 2.5-fold higher eDNA abundance for anchovy than sardine which agree in direction with stock assessments for anchovy and sardine (an approximately 35-fold difference; Kuriyama et al., 2022b,a). However, due to differences in stock definitions and survey areas (the anchovy stock assessment includes southern California waters (32.5*^◦^*–36*^◦^*N) not covered by the eDNA survey) these magnitudes are not directly comparable. Lastly, sardine eDNA also showed elevated concentrations below depths of 150 m – contrary to previous reports indicating only occasional detections at these depths (Zwolinski and Demer, 2024). However, these detections were rare with only 17 positive samples out of 120 in which 15 of those samples contained concentrations lower than 100 copies/L, suggesting that the deep signal likely represents stochastic processes including downward eDNA transport, deposition via predatory feces, carcass decay, transient individuals, or field contamination.

**Figure 5:**
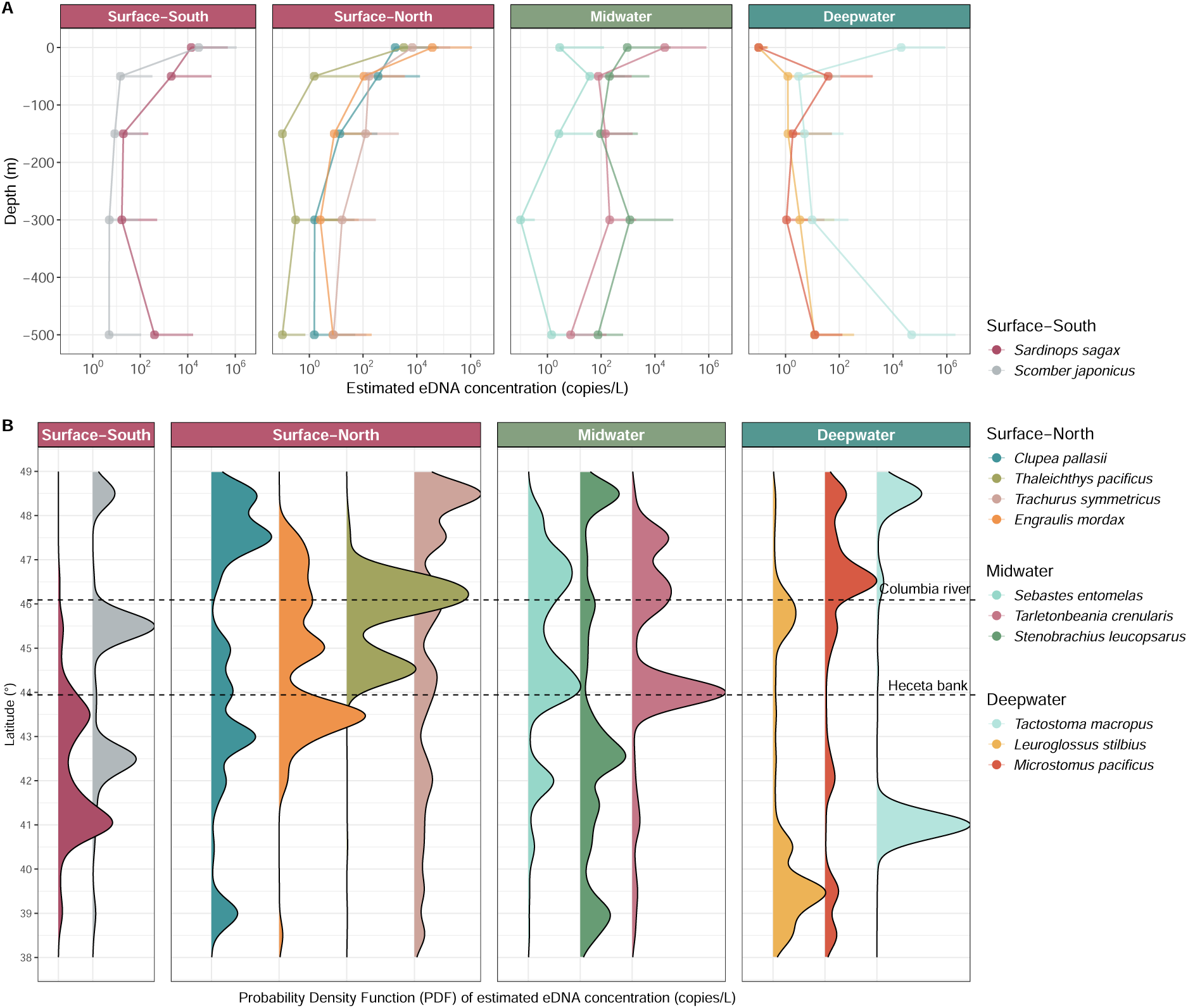
Vertical (A) and latitudinal (B) distribution of eDNA concentrations grouped in ecological groups (surface south, surface north, midwater, and deepwater). Mean eDNA concentration of vertical distribution is shown with dots and 99% upper quantile with ticks (A). Probability density functions (PDF) of estimated eDNA concentrations across latitudinal gradients (B; square root transformed of x-axis to enhance low concentration peaks). Key geographic features, such as the Columbia River and Heceta Bank, are marked to indicate alignment of species distributions.

### Midwater species

Widow rockfish (*Sebastes entomelas*) is common between 100-350m between northern Baja California and southern Alaska, with pelagic larvae and juveniles generally detected in shallower habitats (*<* 150m depth) (Reynolds, 2001; Bosley et al., 2014) including in nearshore kelp forests (Quigley et al., 2024; Laidig et al., 2007). Moreover, because it is a common bycatch species in the Pacific hake (*Merluccius productus*) fishery, concerns about bycatch of widow rockfish can also influence the much larger hake fishery (i.e., it is a “choke” species for the hake fishery; Somers et al., 2018). Despite its commercial importance, no targeted survey effort exists for widow rockfish, in part because bottom trawls do not sample its pelagic habitat well. We find widow rockfish eDNA strongly and nearly exclusively associated with 50m sample depths (Fig. 2 and 5), consistent with the pelagic larvae and juveniles reported in the literature and somewhat shallower than adults (Reynolds, 2001; Bosley et al., 2014; Quigley et al., 2024). Given water samples were taken at night, this pattern potentially reflects the species’ nocturnal feeding schooling behavior (Reynolds, 2001; Bosley et al., 2014).

The eDNA patterns for northern lampfish (*Stenobrachius leucopsarus*) – an abundant myctophid that migrates from 400-700m up to 20-200m at night (Moku et al., 2000; Suntsov and Brodeur, 2008) – are again consistent with the ecology of the species, having notably lower concentrations in surface samples. Likely owing to the diel vertical migration of the northern lampfish, the signal of eDNA for this species is observed across depths.

### Deepwater species

Among deeper-water fishes, eDNA revealed spatial and potentially ontogenetic migration patterns for species that are common, yet under studied, such as the longfin dragonfish (*Tactostoma macropus*). Available literature suggests dragonfish occur roughly between 40*^◦^*N-50*^◦^*N (adults at 300-900m, larvae at 0-60m in summer; Kawaguchi and Moser, 1993) and perform diel vertical migrations to around 100 m (Kawaguchi and Moser, 1993; Willis and Pearcy, 1982). The eDNA concentrations appeared to reflect this activity (Fig. 5), with the predominant surface signal at northern latitudes likely reflecting larvae and juveniles (Fig. 2; near Columbia River discharge), and two deeper hotspots at 300m and 500m separated by about eight degrees of latitude. However, no other quantitative estimates of abundance or detailed distribution patterns for dragonfish exist in the current literature.

Also consistent with known habitats are commercially caught flatfish such as Dover sole (*Microstomus pacificus*), which lives at 200m to 1200m along the Pacific coast (Drazen, 2007; Brodziak and Mikus, 2000; Drazen and Haedrich, 2012); individuals move progressively deeper as they grow (Vetter et al., 1994; Hunter et al., 1990). The eDNA concentration estimates are consistent with this pattern, showing high concentrations mostly in water samples collected close to the seabed (Fig. 2; Ono et al., 2016).

### Species Associations

Pairwise correlations of DNA concentration showed meaningful patterns among species associated with depth (Fig. 3). Surface-associated species were nearly all positively correlated with each other, with some correlations particularly large (Pearson’s *ρ >* 0.50; Fig. 3). Correlations were weak among the six midwater and deepwater species (all *ρ < |*0.40*|*; Fig. 3). Within the surface species, further subdivision was also apparent, with chub mackerel (*Scomber japonicus*) and sardine (*Sardinops sagax*) showing very high correlation (*ρ* = 0.76; Fig. 3) with both more common in the more southern (and somewhat warmer) waters (Fig. 2 and Fig. 5). Concentrations of the remaining surface-dwelling species (northern anchovy, eulachon, jack mackerel, and Pacific herring) had modest correlations with midwater species (Fig. 3) consistent with more northerly distribution (Fig. 5). Together these findings suggest that eDNA is capturing both depth and latitudinal habitat associations. Deep-water species are distinct (Fig. 2 and Fig. 5) and not strongly correlated with those from other zones (Fig. 3). Deeper water tends to feature lower and more diffuse fish eDNA concentrations (Fig. 2 and Fig. 5). Lower overall eDNA concentrations and fewer deep water samples may make similarities among species distributions more difficult to detect at 300m and deeper.

### Species richness and total eDNA accumulation

In addition to providing species-specific patterns, eDNA metrics of abundance from individual species can be combined to reveal important aggregate ecological patterns. For the 12 species considered here, we show how species richness (see SI Appendix, Fig. S2 for defining species presence) from eDNA varied substantially and predictably with depth, with the highest richness observed at the surface (Fig. 4A). In part, this pattern is likely to be an artifact of the focal species included in this study – chosen for their relevance to fisheries management as well as for their frequency in the observations (see Methods) – but the richness-depth gradient also reflects the expectation that habitat and nutrient availability is greatest near the surface (Smith and Brown, 2002; Hickey et al., 2005). Surface waters also host the eggs, larvae, and juvenile stages of many meso- and bathypelagic species (Parnel et al., 2008), some of which we have included here, and hence we both expect and observe higher richness in the upper layer of the water column (Kim and Barth, 2011).

Species richness also corresponds strongly with the influence of Columbia River plume (Fig. 4A), which extends well beyond the coastline (Hickey and Banas, 2003). The nitrate-rich river water interacts with ocean currents and intensifies coastal upwelling, enhancing both primary and secondary productivity (Hickey et al., 2010) and likely influencing the patterns of species richness we observe (Fig. 4A; Tolimieri et al., 2015). The total fish eDNA accumulation (Fig. 4B) suggested the highest aggregation of forage fish DNA near Heceta Bank (44 degrees N; overlapping anchovy and sardine distributions are the two largest contributors to the observed peak of accumulated eDNA concentration). In addition to surface-dwelling species, fish eDNA accumulation revealed high aggregation of two important mesopelagic fishes (*Sebastes entomelas* and *Tarletonbeania crenularis*) – a center of abundance for commercial fisheries (Bosley et al., 2014; Tissot et al., 2008). Although our eDNA-derived richness reflects an index based on a subset of taxa (12 species here) and integrates signals over space and time, its spatial patterns correspond closely with known ecological structures and independently observed biodiversity hotspots.

## Discussion

We report species-specific DNA concentrations smoothed over ten degrees of latitude and across the continental shelf to 500m. We reveal distributional patterns for 12 species ranging from the iconic coastal pelagic species of the northeast Pacific to understudied-but-abundant deepwater species, and our results suggest pathways for the broader use of eDNA in both fisheries and ecosystem management.

### eDNA as Management Tool

The eDNA-derived spatial and vertical patterns revealed here reflect the distributions and ecological associations of a diverse suite of fish species across the California Current. Consistent with previous studies that have demonstrated species-specific relationships between eDNA concentrations and fish biomass – such as those by Salter et al. (2019) for Atlantic cod, Shelton et al. (2022) for hake, Maes et al. (2023) for two commercial flatfishes, and Guri et al. (2024a) for multiple commercial fishes – we find species-level eDNA concentrations align closely with the preferred habitats’ physiological constraints (e.g., temperature tolerance or food availability) described for each species in the available literature. The observed latitudinal differences of eDNA concentration among species – such as those between anchovy and sardine – highlights species’ thermal ranges and largely coincide with other surveys (Zwolinski and Demer, 2024); moreover, eDNA patterns appear to reflect diel vertical migrations (observed in myctophids) and ontogenetic migrations, where eDNA concentrations are consistent with the species’ ontogenetic movement between surface and mid-to deep-water habitats. We also identify aggregations of richness and total eDNA near high-productivity regions of ecological and commercial value. By mapping three-dimensional spatial patterns, eDNA provides information on species distributions, habitat preferences, and ecological processes that is often otherwise unavailable. Using metabarcoding approach jointly with qPCR and mock communities, a single water sample can yield quantitative information on tens to hundreds of species – including those for which traditional fisheries surveys are not currently conducted. However, a few deviations from expected patterns are also apparent – most notably the presence of sardine eDNA at depths exceeding 150 m, where individuals are rarely observed. Such signals likely reflect downward transport processes (e.g., particle sinking, fecal deposition, or carcass decay), or field contaminations rather than true presence at depth.

For well-studied taxa, existing information helps contextualize eDNA signals, providing insights into habitat preferences, seasonal patterns, or temperature tolerances. For poorly studied species, eDNA serves as an exploratory tool, detecting elusive taxa and revealing unexpected habitats. Widow rockfish, for example, appear almost exclusively in pelagic habitats around 50m deep, which are sampled poorly by traditional bottom-trawl surveys (Keller et al., 2017). This species has been of special interest, as its targeted commercial harvest resumed in the late 2010s after a decade of closure (harvest was less than 500 mt annually between 2003 and 2013 but has exceeded 10,000 mt annually since 2018) and no existing survey adequately captures its patterns of abundance (Adams et al., 2019).

### Advantages and Limitations

Determining the abundance and distribution of marine species is challenging as different taxa occupy distinct habitats and respond differently to each type of sampling gear, producing variable detectability and catchability in traditional multispecies surveys (Zhang et al., 2020). Metabarcoding circumvents these limitations by effectively sampling numerous taxa simultaneously with a single “gear,” offering a standardized, scalable, and replicable approach for broad spatial and temporal surveys. The eDNA-derived estimates we provide here have direct relevance for natural-resources management at large scales. For example, indices of abundance have a central role in determining stock status for fisheries applications (i.e., stock assessments in the U.S. under the Magnuson-Stevens Act) as well as informing conservation status (e.g., under the U.S. Endangered Species Act). Thus eDNA-derived abundance indices can fill a variety of existing requirements for management (Baetscher et al., 2025). While these indices are still in their infancy and will require additional technical developments to ensure their proper usage, they are conceptually and practically little different from abundance estimates derived from more traditional ocean survey sampling methods (i.e., net-based or acoustic surveys) (Shelton et al., 2022; Guri et al., 2024a; Maes et al., 2023; Salter et al., 2019). Indeed, eDNA methods may provide some notable improvements because single- and multi-species eDNA approaches avoid tradeoffs involved with physically capturing many species simultaneously using a single net. Molecular surveys capture many species with the same “gear” using methods that are easily replicated in space and time and can be used to generate data for species not typically targeted by existing sampling methods. As agencies worldwide begin to implement eDNA strategies – with the U.S. National Aquatic eDNA Strategy (Kelly et al., 2024) serving as one example on folding eDNA data into federal decision-making – and other large-scale examples point the way to maximizing the benefit of this information-rich datastream for natural-resources management at both continental and global scales, particularly in data-poor regions where traditional monitoring infrastructure is limited.

Our findings illustrate the power of eDNA as a tool for fisheries management, biodiversity conservation, and the study of environmental change. The observable eDNA signal at any given location and time is an integrated product of DNA production and loss across time and space (Gold et al., 2023). In some contexts, it may make sense to aggregate spatially proximate samples to better reflect the combined forces of eDNA production, transport, and decay (Guri et al., 2024a). As in any sampling context, individual samples may over- or underrepresent the true abundance at a given location due to the distance to organisms or eDNA dispersion. These stochastic effects should average out at larger scales and are directly akin to observation variability present in traditional sampling methods (e.g. net or acoustics).

Furthermore, since the production of DNA from the source is continuous and relatively localized in both spatial dimensions (within km; Baetscher et al., 2023) and temporal (within hours; Strickler et al., 2015; Collins et al., 2018; Mächler et al., 2018; Andruszkiewicz Allan et al., 2021), the distribution of eDNA from a given species at any given time will be smoother and more extensive than the distribution of discrete individuals. Such characteristics establish eDNA as a source of continuous, integrated biological information different from traditional methods (i.e., trawling or mark-recapture) which typically rely on the assumption that the fish population is static during the sampling period (Bailey et al., 2014). This feature means that eDNA can reflect biological information that extends beyond the precise sampling location and timeframe. Due to the integrated nature of the eDNA signal and (here) the subsequent model-based smoothing, our findings are best interpreted as representing large-scale ecological patterns rather than localized dynamics.

Vertical eDNA signals in this study largely align with the known depth distributions of most species, and represent a dimension that most traditional survey methods do not capture (i.e., co-located samples at multiple depths). For instance shallow-water species (sardine, anchovy, herring) occur largely at the surface and 50 m depth bins; widow rockfish occurs almost exclusively at 50 m; species found throughout the water column are either diel vertical migrators or species occupying different depths at various life-history stages (Fig. 4B); and bottom-dwelling fishes are detected primarily at the deepest point sampled on any station. However, interpreting these patterns is complicated by multiple factors. First, vertical transport processes - especially downward eDNA flux via particle settling - can obscure biological signals at depth. However, if downward transport were a pervasive mechanism, we would expect to observe similar deep signals for other species; yet widow rockfish eDNA remains tightly constrained around the 150 m depth bin, consistent with its known vertical range and inconsistent with passive settling. Additionally, fecal deposition from predators feeding on sardines in surface waters can transport sardine DNA to deeper layers, where particulate material settles rapidly through the water column. Likewise, the decomposition of recently dead individuals may release DNA that is subsequently transported downward with organic detritus. These mechanisms could explain elevated sardine eDNA in deep layers (*>*150 m; Fig. 4A). This represents a clear case where sardines are known surface dwellers, we suspect high concentration of sardine DNA at depth reflect downward transport processes. However, for other pelagic species that undergo ontogenetic or diel vertical migrations (DVMs), disentangling biological signals becomes more challenging since eDNA cannot identify life stages or pinpoint shedding times; differentiating DNA derived from organismal migration or the passive transport of residual DNA is not currently possible. Despite these limitations, studies have reported that marine communities can be clustered by depth through eDNA surveys (Easson et al., 2020; Jeunen et al., 2020) and mechanistic models from previous studies indicate that eDNA profiles reach quasi-equilibrium within days under stable migration patterns (Andruszkiewicz Allan et al., 2021). This stability supports linking eDNA depth distributions to resident depths (day/night) if migration consistency is assumed. In summary, the overall observed depth-specific pattern of eDNA signals here align with literature reports and known species depth preferences; however, detailed interpretations require caution and should consider the confounding effects of vertical eDNA transport.

Direct comparison of our eDNA-derived patterns against independent observations of fish biomass falls beyond the scope of this study, particularly given the lack of observational or repository data for many species included here. Consequently, we cannot quantify the method’s uncertainty relative to “true” biological signals, since a comprehensive uncertainty assessment would require cross-validation with alternative observation techniques—data that are currently unavailable and themselves subject to different biases. However, an element of our method’s uncertainty can be indicated through the analysis of smoothing residuals, which compare spatially smoothed maps (Fig. 2 and SI Appendix, Fig. S1) with sample-specific eDNA estimates derived from the joint model (SI Appendix, Fig. S5). While spatial smoothing assumes a continuous surface between sampling locations, each eDNA sample is an independent observation, allowing the residuals to highlight localized deviations. Therefore, the absolute difference between the observed values and the smoothed predictions—the smoothing residuals—can be an indicator of the observation variability.

### Integrating DNA with traditional surveys

Given that our sampling area spans more than 1,200 km north to south and tens of km east to west – significantly exceeding the scale of potential eDNA transport and decay – the observed spatial patterns we report are likely to reflect genuine differences in fish communities rather than artifacts of eDNA horizontal transport dynamics (Guri et al., 2024b).

Effective management of natural resources at large scales requires that we maximize the information content of existing sampling regimes. Here, we follow Shelton et al. (2022), leveraging the same water samples to provide information on an additional 12 species of ecosystem and management importance. Elsewhere, a subset of these samples has yielded information on the distribution of marine mammals (Valdivia-Carrillo et al., 2025, in revision). The repeated use of a single set of samples highlights an important strength of eDNA samples; they allow researchers to revisit and re-sequence the genetic material in archived samples as new research questions arise or attention shifts to different taxa. The use of these samples, from a targeted fisheries survey for hake, highlights the efficiency for which eDNA can effectively provide survey data for tens or hundreds of additional species for which traditional surveys do not exist.

This capacity for retrospective analysis allows historical samples to be re-examined at any point, insofar as the sample is not exhausted, thereby offering opportunities to address emerging priorities and to reconstruct past ecological conditions. For example, the ability to retrospectively pinpoint the origins and pathways of invasive species can inform targeted interventions, helping managers understand not only where an incursion began but also how it spread (Gilbey et al., 2021). Similarly, the capacity to trace the initial onset of harmful algal blooms or the earliest presence of pathogenic microbes enables researchers and policymakers to identify the underlying drivers of these events and mitigate future outbreaks (Shaw et al., 2019).

Moreover, integrating retrospective eDNA analyses a changing climate can enrich predictive frameworks by providing historically grounded baselines of community composition, thereby enhancing our understanding of ecosystem responses to environmental shifts. For example, Díaz et al. (2020) demonstrated how archived samples could be leveraged to reconstruct past fish communities using eDNA, thereby complementing traditional monitoring methods and further extend the utility of archived material. Such retrospective applications deepen and extend the value of every research expedition, ensuring that once-collected samples continue to yield valuable ecological insights over time. We therefore see eDNA as an important data source that maximizes the information value of existing research cruises, rather than a means of replacing those cruises themselves.

We have shown an example of the potential value of multi-species eDNA analysis for fisheries and other natural-resource management questions (Ledger et al., 2024; Stoeckle et al., 2024), however, molecular patterns alone are unlikely to drive quantitative natural-resources decisions, in part because eDNA generally does not provide auxiliary information important for managing populations (e.g., information on size- or age structure, fecundity, sex, or disease). Relating observed eDNA concentrations to species mass requires data external to the eDNA signal (Guri et al., 2024a); and in particular, requires that we understand proportionalities between eDNA concentration and biomass that may vary among species. We therefore see eDNA analysis as strengthening and extending existing surveys, with different data streams complementing one another.

## Author Contributions

AOS, KMN, RPK, MS, and KP designed the research; MRS, AR-L and AW did the laboratory work; PB, RPK, and GG did the bioinformatics; GG, OL, RPK, and AOS analyzed the data; GG, OL, RPK, and AOS interpreted the results; All authors provided edits to the manuscript

## Author Declaration

The authors declare no competing interest.

## Acknowledgment

We thank Linda Park for her foresight and perseverance, Katherine Maslenikov (University of Washington Burke Museum Ichthyology Collection) for providing tissue samples for the mock community analysis, and Olivia Scott for validation and Sanger sequencing of the mock communities. This material is based upon research supported by the Office of Naval Research under Award Number (N00014-22-1-2719) and support from the National Marine Fisheries Service’s Genomic Strategic Initiative. The authors gratefully acknowledge the survey expertise and support of NOAA NWFSC’s Pacific hake survey team and the personnel of the NOAA Ship Bell M. Shimada for supporting the collection of eDNA samples on the west coast hake survey 2019 - 2023. Any use of trade, firm, or product names is for descriptive purposes only and does not imply endorsement by the U.S. Government.

## References

1. Adams, G. D., Kapur, M., McQuaw, K., Thurner, S., Hamel, O., Stephens, A., and Wetzel, C. (2019). Stock Assessment Update: Status of Widow Rockfish (*Sebastes Entomelas*) along the U.S. West Coast in 2019. Stock Assessment Update, Pacific Fishery Management Council, Portland, Oregon.

2. Allan, E. A., Kelly, R. P., D’Agnese, E. R., Garber-Yonts, M. N., Shaffer, M. R., Gold, Z. J., and Shelton, A. O. (2023). Quantifying impacts of an environmental intervention using environmental DNA. Ecological Applications, 33(8):e2914.

3. Anderson, S. C., Ward, E. J., English, P. A., Barnett, L. A. K., and Thorson, J. T. (2022). sdmTMB: An R Package for Fast, Flexible, and User-Friendly Generalized Linear Mixed Effects Models with Spatial and Spatiotemporal Random Fields. *bioRxiv*.

4. Andres, K. J., Lodge, D. M., and Andŕes, J. (2023). Environmental DNA reveals the genetic diversity and population structure of an invasive species in the Laurentian Great Lakes. Proceedings of the National Academy of Sciences, 120(37):e2307345120.

5. Andruszkiewicz Allan, E., Zhang, W. G. C., Lavery, A., and F. Govindarajan, A. (2021). Environmental DNA shedding and decay rates from diverse animal forms and thermal regimes. Environmental DNA, 3(2):492–514.

6. Baetscher, D. S., Nuetzel, H. M., and Garza, J. C. (2023). Highly accurate species identification of Eastern Pacific rockfishes (Sebastes spp.) with high-throughput DNA sequencing. Conservation Genetics, 24(5):563–574.

7. Baetscher, D. S., Omori, K. L., Goethel, D. R., Shelton, A. O., Berger, A. M., Ledger, K. J., Nichols, K. M., and Larson, W. A. (2025). The Pragmatic Sceptic: A Practical Approach for Integrating Environmental DNA Into Marine Stock Assessment and Fisheries Management. Fish and Fisheries, page faf.70001.

8. Baetscher, D. S., Pochardt, M. R., Barry, P. D., and Larson, W. A. (2024). Tide impacts the dispersion of eDNA from nearshore net pens in a dynamic high-latitude marine environment. Environmental DNA, 6(2):e533.

9. Bailey, L. L., MacKenzie, D. I., and Nichols, J. D. (2014). Advances and applications of occupancy models. Methods in Ecology and Evolution, 5(12):1269–1279.

10. Bosley, K. L., Miller, T. W., Brodeur, R. D., Bosley, K. M., Van Gaest, A., and Elz, A. (2014). Feeding ecology of juvenile rockfishes off Oregon and Washington based on stomach content and stable isotope analyses. Marine Biology, 161(10):2381–2393.

11. Brodziak, J. and Mikus, R. (2000). Variation in life history parameters of Dover sole, *Microstomus pacificus*, off the coasts of Washington, Oregon, and northern California. Fisheries Bulletin, 98(4):661–673.

12. Callahan, B. J., McMurdie, P. J., Rosen, M. J., Han, A. W., Johnson, A. J. A., and Holmes, S. P. (2016). DADA2: High-resolution sample inference from Illumina amplicon data. Nature Methods, 13(7):581–583.

13. Collins, R. A., Wangensteen, O. S., O’Gorman, E. J., Mariani, S., Sims, D. W., and Genner, M. J. (2018). Persistence of environmental DNA in marine systems. Communications Biology, 1(1):185.

14. Cornelis, I., De Backer, A., Maes, S., Vanhollebeke, J., Brys, R., Ruttink, T., Hostens, K., and Derycke, S. (2024). Environmental DNA for monitoring the impact of offshore wind farms on fish and invertebrate community structures. Environmental DNA, 6(4):e575.

15. de Blois, S. (2020). The 2019 Joint U.S.-Canada Integrated Ecosystem and Pacific Hake Acoustic-Trawl Survey: Cruise Report SH-19-06. NMFS-NWFSC-PR-2020-03, U.S. Department of Commerce, NOAA Processed Report.

16. Díaz, C., Wege, F.-F., Tang, C. Q., Crampton-Platt, A., Rüdel, H., Eilebrecht, E., and Koschorreck, J. (2020). Aquatic suspended particulate matter as source of eDNA for fish metabarcoding. Scientific Reports, 10(1):14352.

17. DiBattista, J. D., Fowler, A. M., Riley, I. J., Reader, S., Hay, A., Parkinson, K., and Hobbs, J.-P. A. (2022). The use of environmental DNA to monitor impacted coastal estuaries. Marine Pollution Bulletin, 181:113860.

18. Drazen, C. J. (2007). Depth related trends in proximate composition of demersal fishes in the eastern North Pacific. Deep Sea Research Part I: Oceanographic Research Papers, 54(2):203–219.

19. Drazen, J. C. and Haedrich, R. L. (2012). A continuum of life histories in deep-sea demersal fishes. Deep Sea Research Part I: Oceanographic Research Papers, 61:34–42.

20. Easson, C. G., Boswell, K. M., Tucker, N., Warren, J. D., and Lopez, J. V. (2020). Combined eDNA and Acoustic Analysis Reflects Diel Vertical Migration of Mixed Consortia in the Gulf of Mexico. Frontiers in Marine Science, 7:552.

21. Gilbey, J., Carvalho, G., Castilho, R., Coscia, I., Coulson, M. W., Dahle, G., Derycke, S., Francisco, S. M., Helyar, S. J., Johansen, T., Junge, C., Layton, K. K., Martinsohn, J., Matejusova, I., Robalo, J. I., Rodŕıguez-Ezpeleta, N., Silva, G., Strammer, I., Vasemägi, A., and Volckaert, F. A. (2021). Life in a drop: Sampling environmental DNA for marine fishery management and ecosystem monitoring. Marine Policy, 124:104331.

22. Gloor, G. B., Macklaim, J. M., Pawlowsky-Glahn, V., and Egozcue, J. J. (2017). Microbiome datasets are compositional: And this is not optional. Frontiers in Microbiology, 8:2224.

23. Gold, Z. J., Shelton, A. O., Casendino, H. R., Duprey, J., Gallego, R., Van Cise, A., Fisher, M., Jensen, A. J., D’Agnese, E., Andruszkiewicz Allan, E., Ramón-Laca, A., Garber-Yonts, M., Labare, M., Parsons, K. M., and Kelly, R. P. (2023). Signal and noise in metabarcoding data. PLOS ONE, 18(5):e0285674.

24. Guri, G., Shelton, A. O., Kelly, R. P., Yoccoz, N., Johansen, T., Præbel, K., Hanebrekke, T., Ray, J. L., Fall, J., and Westgaard, J.-I. (2024a). Predicting trawl catches using environmental DNA. ICES Journal of Marine Science, 81(8):1536–1548.

25. Guri, G., Westgaard, J.-I., Yoccoz, N., Wangensteen, O. S., Præbel, K., Ray, J. L., Kelly, R. P., Shelton, A. O., Hanebrekke, T., and Johansen, T. (2024b). Maximizing sampling efficiency to detect differences in fish community composition using environmental DNA metabarcoding in subarctic fjords. Environmental DNA, 6(1):e409.

26. Hickey, B., Geier, S., Kachel, N., and MacFadyen, A. (2005). A bi-directional river plume: The Columbia in summer. Continental Shelf Research, 25(14):1631–1656.

27. Hickey, B. M. and Banas, N. S. (2003). Oceanography of the U.S. Pacific Northwest Coastal Ocean and estuaries with application to coastal ecology. Estuaries, 26(4):1010–1031.

28. Hickey, B. M., Kudela, R. M., Nash, J. D., Bruland, K. W., Peterson, W. T., MacCready, P., Lessard, E. J., Jay, D. A., Banas, N. S., Baptista, A. M., Dever, E. P., Kosro, P. M., Kilcher, L. K., Horner-Devine, A. R., Zaron, E. D., McCabe, R. M., Peterson, J. O., Orton, P. M., Pan, J., and Lohan, M. C. (2010). River Influences on Shelf Ecosystems: Introduction and synthesis. Journal of Geophysical Research: Oceans, 115:C00B17.

29. Hunter, J. R., Butler, J. L., Kimbrell, C., and Lynn, E. (1990). Bathymetric patterns in size, age, sexual maturity, water content, and caloric density of Dover sole, Microstomus pacificus. California Cooperative Oceanic Fisheries Investigations Reports, 31:132–144.

30. Jeunen, G., Lamare, M. D., Knapp, M., Spencer, H. G., Taylor, H. R., Stat, M., Bunce, M., and Gemmell, N. J. (2020). Water stratification in the marine biome restricts vertical environmental DNA (eDNA) signal dispersal. Environmental DNA, 2(1):99–111.

31. Jo, T., Murakami, H., Yamamoto, S., Masuda, R., and Minamoto, T. (2019). Effect of water temperature and fish biomass on environmental DNA shedding, degradation, and size distribution. Ecology and Evolution, 9(3):1135–1146.

32. Johnson, K. F., Edwards, A. M., Berger, A. M., Grandin, C. J., and Wetzel, C. R. (2025). Status of the pacific hake (whiting) stock in U.S. and Canadian waters in 2025. Technical report, Joint Technical Committee of the U.S. and Canada Pacific Hake/Whiting Agreement; National Marine Fisheries Service; Fisheries and Oceans Canada, Seattle, WA, USA and Nanaimo, B.C., Canada.

33. Kaplan, I. C., Francis, T. B., Punt, A. E., Koehn, L. E., Curchitser, E., Hurtado-Ferro, F., Johnson, K. F., Lluch-Cota, S. E., Sydeman, W. J., Essington, T. E., Taylor, N., Holsman, K., MacCall, A. D., and Levin, P. S. (2019). A multi-model approach to understanding the role of Pacific sardine in the California Current food web. Marine Ecology Progress Series, 617-618:307–321.

34. Kawaguchi, K. and Moser, H. G. (1993). Development and distribution of the early life history stages of the mesopelagic fish *Tactostoma macropus* (stomiidae) in the transitional waters of the north pacific. Japanese Journal of Ichthyology, 40(2):161–172.

35. Keller, A. A., Wallace, J. R., and Methot, R. D. (2017). The Northwest Fisheries Science Center’s West Coast Groundfish Bottom Trawl Survey: History, Design, and Description. NMFS-NWFSC-136, U.S. Department of Commerce, NOAA Technical Memorandum.

36. Kelly, R. P., Lodge, D. M., Lee, K. N., Theroux, S., Sepulveda, A. J., Scholin, C. A., Craine, J. M., Andruszkiewicz Allan, E., Nichols, K. M., Parsons, K. M., Goodwin, K. D., Gold, Z., Chavez, F. P., Noble, R. T., Abbott, C. L., Baerwald, M. R., Naaum, A. M., Thielen, P. M., Simons, A. L., Jerde, C. L., Duda, J. J., Hunter, M. E., Hagan, J. A., Meyer, R. S., Steele, J. A., Stoeckle, M. Y., Bik, H. M., Meyer, C. P., Stein, E., James, K. E., Thomas, A. C., Demir-Hilton, E., Timmers, M. A., Griffith, J. F., Weise, M. J., and Weisberg, S. B. (2024). Toward a national eDNA strategy for the United States. Environmental DNA, 6(1):e432.

37. Kelly, R. P., Shelton, A. O., and Gallego, R. (2019). Understanding PCR processes to draw meaningful conclusions from environmental DNA studies. Scientific Reports, 9(1):12133.

38. Kim, S. and Barth, J. A. (2011). Connectivity and larval dispersal along the Oregon coast estimated by numerical simulations. Journal of Geophysical Research, 116(C6):C06002.

39. Kuriyama, P. T., Zwolinski, J. P., and Hill, K. T. (2022a). Update Assessment of the Pacific sardine resource in 2022 for U.S. management in 2022-2023. Technical Report NMFS-SWFSC-662, U.S. Department of Commerce, NOAA Technical Memorandum.

40. Kuriyama, P. T., Zwolinski, J. P., Teo, S. L., and Hill, K. T. (2022b). Assessment of the northern anchovy (*Engraulis mordax*) central subpopulation in 2021 for U.S. management. Technical Report NMFS-SWFSC-665, U.S. Department of Commerce, NOAA Technical Memorandum.

41. Laidig, T. E., Chess, J. R., and Howard, D. F. (2007). Relationship between abundance of juvenile rockfishes (Sebastes spp.) and environmental variables documented off northern California and potential mechanisms for the covariation. Fisheries Bulletin, 105(39-48):39–48.

42. Langlois, V. S., Allison, M. J., Bergman, L. C., To, T. A., and Helbing, C. C. (2021). The need for robust qPCR-based eDNA detection assays in environmental monitoring and species inventories. Environmental DNA, 3(3):519–527.

43. Ledger, K. J., Everett, M., Nichols, K. M., Larson, W. A., and Baetscher, D. S. (2025). Development and Validation of an eDNA Primer Set for Rockfishes in the Alaska Current Provides a New Tool for Fisheries Management. Environmental DNA, 7(4):e70176.

44. Ledger, K. J., Hicks, M. B. R., Hurst, T. P., Larson, W., and Baetscher, D. S. (2024). Validation of environmental DNA for estimating proportional and absolute biomass. Environmental DNA, 6(5):e70030.

45. Litz, M. N. C., Heppell, S. A., Emmett, R. L., and Brodeur, R. D. (2008). Ecology and distribution of the northern subpopulation of northern anchovy (*Engraulis mordax*) off the U.S. West Coast. California Cooperative Oceanic Fisheries Investigations Reports, 49:167–182.

46. Liu, O. R., Ward, E. J., Anderson, S. C., Andrews, K. S., Barnett, L. A., Brodie, S., Carroll, G., Fiechter, J., Haltuch, M. A., Harvey, C. J., Hazen, E. L., Hernvann, P.-Y., Jacox, M., Kaplan, I. C., Matson, S., Norman, K., Buil, M. P., Selden, R. L., Shelton, A., and Samhouri, J. F. (2023). Species redistribution creates unequal outcomes for multispecies fisheries under projected climate change. Science Advances, 9(33):eadg5468.

47. Mächler, E., Osathanunkul, M., and Altermatt, F. (2018). Shedding light on eDNA: Neither natural levels of UV radiation nor the presence of a filter feeder affect eDNA-based detection of aquatic organisms. PLOS ONE, 13(4):e0195529.

48. Maes, S. M., Desmet, S., Brys, R., Sys, K., Ruttink, T., Maes, S., Hostens, K., Vansteenbrugge, L., and Derycke, S. (2023). Detection and quantification of two commercial flatfishes *Solea solea* and *Pleuronectes platessa* in the North Sea using environmental DNA. Environmental DNA, 6(1):e426.

49. Martin, M. (2011). Cutadapt removes adapter sequences from high-throughput sequencing reads. EMB-net.journal, 17(1):10.

50. McLaren, M. R., Willis, A. D., and Callahan, B. J. (2019). Consistent and correctable bias in metagenomic sequencing experiments. eLife, 8:e46923.

51. Miya, M., Sato, Y., Fukunaga, T., Sado, T., Poulsen, J. Y., Sato, K., Minamoto, T., Yamamoto, S., Yamanaka, H., Araki, H., Kondoh, M., and Iwasaki, W. (2015). MiFish, a set of universal PCR primers for metabarcoding environmental DNA from fishes: Detection of more than 230 subtropical marine species. Royal Society Open Science, 2:150088.

52. Moku, M., Kawaguchi, K., Watanabe, H., and Ohno, A. (2000). Feeding habits of three dominant myctophid fishes, *Diaphus theta*, *Stenobrachius leucopsarus* and *S. nannochir*, in the subarctic and transitional waters of the Western North Pacific. Marine Ecology Progress Series, 207:129–140.

53. Muenzel, D., Bani, A., De Brauwer, M., Stewart, E., Djakiman, C., Halwi, Purnama R., Yusuf, S., Santoso, P., Hukom, F. D., Struebig, M., Jompa, J., Limmon, G., Dumbrell, A., and Beger, M. (2024). Combining environmental DNA and visual surveys can inform conservation planning for coral reefs. Proceedings of the National Academy of Sciences, 121(17):e2307214121.

54. Ono, K., Shelton, A. O., Ward, E. J., Thorson, J. T., Feist, B. E., and Hilborn, R. (2016). Space-time investigation of the effects of fishing on fish populations. Ecological Applications, 26(2):392–406.

55. Parnel, M. M., Emmett, R. L., and Brodeur, R. D. (2008). Ichthyoplankton community in the Columbia River plume off Oregon: Effects of fluctuating oceanographic conditions. Fishery Bulletin, 106(2):161–173.

56. Pikitch, E. K., Boersma, P. D., Boyd, I. L., Conover, D. O., Cury, P. M., Essington, T. E., Heppell, S. S., Houde, E. D., Mangel, M., Pauly, D., Plaganyi-Lloyd, E., Sainsbury, K., and Steneck, R. S. (2012). Little fish, big impact: Managing a crucial link in ocean food webs. In Proceedings of the Lenfest Ocean Program. Report from the Lenfest Forage Fish Task Force.

57. Pont, D. (2024). Predicting downstream transport distance of fish eDNA in lotic environments. Molecular Ecology Resources, 24(4):e13934.

58. Quigley, L. A., Franks, P. J. S., Thompson, A. R., Field, J. C., and Santora, J. A. (2024). Quantifying the fates and retention of larval rockfish through Lagrangian analyses. Marine Ecology Progress Series, 749:109–125.

59. R Core Team (2024). R: A Language and Environment for Statistical Computing.

60. Ramón-Laca, A., Wells, A., and Park, L. (2021). A workflow for the relative quantification of multiple fish species from oceanic water samples using environmental DNA (eDNA) to support large-scale fishery surveys. PLOS ONE, 16(9):e0257773.

61. Reynolds, J. A. (2001). Habitat Associations and Distribution of Widow Rockfish, Sebastes entomelas, with Implications for Marine Reserve Design. PhD thesis, University of California, Berkeley.

62. Ruppert, K. M., Davis, D. R., Rahman, M. S., and Kline, R. J. (2022). Development and assessment of an environmental DNA (eDNA) assay for a cryptic Siren (Amphibia: Sirenidae). Environmental Advances, 7:100163.

63. Salter, I., Joensen, M., Kristiansen, R., Steingrund, P., and Vestergaard, P. (2019). Environmental DNA concentrations are correlated with regional biomass of Atlantic cod in oceanic waters. Communications Biology, 2(1):1–9.

64. Shaw, J. L. A., Weyrich, L. S., Hallegraeff, G., and Cooper, A. (2019). Retrospective eDNA assessment of potentially harmful algae in historical ship ballast tank and marine port sediments. Molecular Ecology, 28(10):2476–2485.

65. Shelton, A. O., Gold, Z. J., Jensen, A. J., D’Agnese, E., Andruszkiewicz Allan, E., Van Cise, A., Gallego, R., Ramón-Laca, A., Garber-Yonts, M., Parsons, K., and Kelly, R. P. (2023). Toward quantitative metabarcoding. Ecology, 104(2):e3906.

66. Shelton, A. O., Kelly, R. P., O’Donnell, J. L., Park, L., Schwenke, P., Greene, C., Henderson, R. A., and Beamer, E. M. (2019). Environmental DNA provides quantitative estimates of a threatened salmon species. Biological Conservation, 237:383–391.

67. Shelton, A. O., Ramón-Laca, A., Wells, A., Clemons, J., Chu, D., Feist, B. E., Kelly, R. P., Parker-Stetter, S. L., Thomas, R., Nichols, K. M., and Park, L. (2022). Environmental DNA provides quantitative estimates of Pacific hake abundance and distribution in the open ocean. Proceedings of the Royal Society B: Biological Sciences, 289(1971):20212613.

68. Smith, K. F. and Brown, J. H. (2002). Patterns of diversity, depth range and body size among pelagic fishes along a gradient of depth. Global Ecology and Biogeography, 11(4):313–322.

69. Somers, K. A., Pfeiffer, L., Miller, S., and Morrison, W. (2018). Using incentives to reduce bycatch and discarding: Results under the West Coast catch share program. Coastal Management, 46(6):621–637.

70. Stan Development Team (2023). Stan Modeling Language Users Guide and Reference Manual.

71. Steinbeck, J. (1945). Cannery Row. Viking Press.

72. Stoeckle, M. Y., Ausubel, J. H., and Coogan, M. (2022). 12S gene metabarcoding with DNA standard quantifies marine bony fish environmental DNA , identifies threshold for reproducible detection, and overcomes distortion due to amplification of non-fish DNA. Environmental DNA, 6(1):e376.

73. Stoeckle, M. Y., Ausubel, J. H., Hinks, G., and VanMorter, S. M. (2024). A potential tool for marine biogeography: eDNA-dominant fish species differ among coastal habitats and by season concordant with gear-based assessments. PLOS ONE, 19(11):e0313170.

74. Strickler, K. M., Fremier, A. K., and Goldberg, C. S. (2015). Quantifying effects of UV-B, temperature, and pH on eDNA degradation in aquatic microcosms. Biological Conservation, 183:85–92.

75. Suntsov, A. V. and Brodeur, R. D. (2008). Trophic ecology of three dominant myctophid species in the northern California Current region. Marine Ecology Progress Series, 373:81–96.

76. Tissot, B. N., Wakefield, W. W., Hixon, M. A., and Clemons, J. (2008). Twenty years of fish-habitat studies on heceta bank, oregon. In Reynolds, J. and Greene, H., editors, Marine Habitat Mapping Technology for Alaska. Alaska Sea Grant for North Pacific Research Board.

77. Tolimieri, N., Shelton, A. O., Feist, B. E., and Simon, V. (2015). Can we increase our confidence about the locations of biodiversity ‘hotspots’ by using multiple diversity indices? Ecosphere, 6(12):1–13.

78. Valdivia-Carrillo, T., Shaffer, M., Parsons, K., Jacobson, E. K., Shelton, A. O., Im, A., Nichols, K. M., Wells, A., Ramón-Laca, A., Kelly, R. P., and Van Cise, A. (2025). Leveraging metabarcoding and generalized additive models to describe cetacean edna distribution along the washington state coast using opportunistic samples. In Revision.

79. Veilleux, H. D., Misutka, M. D., and Glover, C. N. (2021). Environmental DNA and environmental RNA: Current and prospective applications for biological monitoring. Science of The Total Environment, 782:146891.

80. Vetter, R. D., Lynn, E. A., de la Garza, M., and Costa, A. S. V. (1994). Depth zonation and metabolic adaptation in Dover sole, Microstomus pacificus, and other deep-living flatfishes: Factors that affect the sole. Marine Biology, 120(1):145–159.

81. Willis, J. M. and Pearcy, W. G. (1982). Vertical distribution and migration of fishes of the lower mesopelagic zone off Oregon. Marine Biology, 70(1):87–98.

82. Xiong, J., MacCready, P., Brasseale, E., Andruszkiewicz Allan, E., Ramón-Laca, A., Parsons, K. M., Shaffer, M., and Kelly, R. P. (2025). Advective Transport Drives Environmental DNA Dispersal in an Estuary. Environmental Science & Technology, 59(15):7506–7516.

83. Zhang, S., Lu, Q., Wang, Y., Wang, X., Zhao, J., and Yao, M. (2020). Assessment of fish communities using environmental DNA: Effect of spatial sampling design in lentic systems of different sizes. Molecular Ecology Resources, 20(1):242–255.

84. Zwolinski, J. P. and Demer, D. A. (2024). An updated model of potential habitat for northern stock Pacific sardine (*Sardinops sagax*) and its use for attributing survey observations and fishery landings. Fisheries Oceanography, 33(3):e12664.

