## Supplementary material for "Quantitative, Multispecies Monitoring at a Continental Scale": SI Appendix

October 28, 2025

1 School of Marine and Environmental Affairs, University of Washington, Seattle, Washington, USA
2 Northwest Fisheries Science Center, NOAA Fisheries, Seattle, Washington, USA
3 Museo Nacional de Ciencias Naturales, Madrid, Spain

### Extended Methods

#### qPCR

Environmental samples were analyzed in triplicate using the multiplexed assay targeting Pacific hake, Eulachon and Pacific lamprey as described in Ramón-Laca et al. (2021) on a QuanStudio 6 (Applied Biosystems). Only Pacific hake qPCR data was used in this study due to the remaining qPCR targets having little positive amplification.

The samples underwent real-time thermocycler protocol including an initial denaturation step at 95°C 10 min followed by 45 cycles of 15 s at 95°C and 1 min at 60°C. All samples were run in 10 $\mu$L volume consisting of 1 x TaqPath ProAmp Multiplex Master Mix, 0.9 $\mu$M of each primer (forward: AAATGTTTAAACTAGAGCCGAATAGC and reverse: TCGTGGAGTCAAAGTGGGGTAGA), 0.2 $\mu$M of probe (6FAM-CACTCGAGGCCACGAAGTACAATT-(MGB)NFQ), and 2 $\mu$L of DNA template or water for the non-template control. All reactions included 100 copies of internal positive control (IPC) to detect PCR inhibition. Any IPC delay exceeding 0.5 cycles in the non-template control was considered inhibition and inhibited samples were diluted and re-analyzed.

Alongside environmental samples standard samples were analyzed constructed from a 130 bp synthetic DNA fragment (gBlock; IDT) representing the 12S region of Pacific hake (*Merluccius productus*), which encompasses the 101 bp 12S qPCR target (sequence information available in Ramón-Laca et al. (2021)). The synthetic DNA fragment was diluted to create a series of standards with final concentrations ranging from $10^0$ to $10^5$ copies/$\mu$L.

#### Metabarcoding

In total, 568 samples, including 554 environmental samples, 7 PCR blanks and 7 positive controls (holding only kangaroo DNA) were amplified using MiFish-U universal primers (Miya et al., 2015) with Illumina tails (forward TCGTCGGCAGCGTCAGATGTGTATAAGAGACAGGCCGGTAAAACTCGTGCCAGC; reverse GTCT- CGTGGGCTCGGAGATGTGTATAAGAGACAGCATAGTGGGGTATCTAATCCCAGTTTG) by using a two-step PCR protocol. The DNA was amplified in the first PCR reaction (PCR1) in a 20 µL reaction consisting of: 10 µL of Phusion Master Mix (2X), 0.4 µL of the forward primer (10 µM); 0.4 µL of the reverse primer (10 µM), 0.6 µL of 100% DMSO, 0.5 µL of rAlbumin (20 µg/µL), 4.4 µL of nuclease-free water and 2 µL of DNA template. Reactions were run with the following cycling conditions: an initial denaturation of

98°C for 30 sec; followed by 35 cycles of 98°C for 10 sec, 60°C for 30 sec, and of 72°C for 3 sec; with a final extension of 72°C for 10 min and hold at 4°C.

PCR product was cleaned using Ampure Beads (1.2x) and then indexed in a second PCR reaction (PCR2), with the following recipe: 12.5 $\mu$L of KAPA HiFi HotStart ReadyMix (Roche Diagnostics), 1.25 $\mu$L of an index from IDT for Illumina DNA/RNA UD Indexes (Sets A-D), 5 $\mu$L of PCR1 product, and 6.25 $\mu$L of nuclease free water. Cycling conditions included an initial denaturation of 95°C for 5 min; 8 cycles of: 98°C for 20 sec, 56°C for 30 sec, 72°C for 1 min; and a final extension of 72°C for 5 min. Resulting products were visualized on a 2% agarose gel and quantified using Quant-iT™ dsDNA Assay Kit (Thermo Fisher Scientific, USA) with Fluoroskan™ Microplate Fluorometer (Thermo Fisher Scientific, USA). Indexed products were normalized by concentration, pooled into libraries for sequencing, and then size- selected to extract only the target fish band using the E-Gel™ SizeSelect™ II Agarose Gels (Thermo Fisher Scientific, USA). Subsequently the libraries were sequenced on Illumina MiSeq platform using the v3 600 cycle kit.

#### Mock community

We constructed four pairs of mock communities with even and skewed species total genomic DNA using Qubit HS assay quantification method. We then quantified the concentration of the 12S rRNA gene using MarVer1 primers (forward: CGTGCCAGCCACCGCG; reverse: GGGTATCTAATCCYAGTTTG (Valsecchi et al., 2020)), which perfectly matched the template to give unbiased estimates of concentration, using ddPCR (Bio-Rad, Inc., QX200 Droplet Digital PCR system). Each tissue of each species was quantified in a 22 $\mu$L reaction consisted of 2 $\mu$L of DNA template from genomic DNA , 11 $\mu$L of ddPCR EvaGreen (Bio-Rad), 0.22 $\mu$L of each forward and reverse primers (10 uM), and 0.56 $\mu$L of nuclease free water (see SI Appendix Table S.1 for DNA quantification results). The thermocycler reactions were run in C1000 Touch Thermal Cycler with 96-Deep Well Reaction Module (Bio-Rad) using the PCR program as follows: 2 min at 50°C for enzyme activation, 2 min at 95°C for initial denaturation, and 40 cycles of denaturation for 1 sec at 95°C and primer annealing and elongation for 30 sec min at 60°C, with a ramp rate of 2°C per s and heldat 4°C until droplets were read. Droplets were determined to be positive after drawing a threshold based on NTCs.

#### Bioinformatics

Starting with Illumina-demultiplexed sequencing files, we first removed primer and adapter sequences using Cutadapt v4.9, applying the default mismatch tolerance parameter (-e 0.1; Martin, 2011). Next, we performed quality filtering using the filterAndTrim function from the DADA2 R package, retaining its default parameters while specifying the optional truncLen argument to match the minimum MiFish amplicon length. After filtering, we denoised the sequences using DADA2 with default settings (Callahan et al., 2016), which resulted in a list of Amplicon Sequence Variants (ASVs) across all samples. Each resulting ASV was then assigned a unique 28-character hash code using the sha-1 algorithm with the R function map_chr. This allowed us to create a common database of ASVs by hash, enabling us to track which sequences have already been annotated for consistency and to avoid redundant computational efforts.

For taxonomic assignment of ASVs, we used standalone BLAST (Altschul et al., 1990) to compare each unique ASV against the NCBI nr eukaryote database as of January 2025. BLAST arguments were as follows: ”-word_size 30 -evalue ”1e-30” -max_target_seqs 50”. To streamline annotation, we also used the ”-negative_taxids” argument, including a list of target organisms closely related to those present in the sampled region, but known to be absent. The complete list of taxids that were excluded is available alongside the pipeline code. Finally, we performed a least common ancestor (LCA) analysis on the top BLAST hits using TaxonKit v0.17 (Shen and Ren, 2021). Sequences with a 100 percent identity match to any database sequence were prioritized for LCA. In the absence of perfect matches, we conducted LCA analysis on the top hits with identity scores above 99.3, , 98, or 96 percent respectively. In the absence of any hits above 96 identity, taxonomic assignment was truncated. Finally, for mock communities were identify of species was known, we manually curated annotations before analysis.

#### Statistical model

##### Overview

We developed a joint model for modeling DNA concentration in the coastal ocean using single-species qPCR data Pacific hake and metabarcoding data from the 12S MiFish Universal primer. This work builds off previous single-species eDNA models constructed for Pacific hake (Shelton et al., 2022) and models for joining qPCR and metabarcoding data (Guri et al., 2024). Here qPCR observations provide information about DNA concentration for one species (Pacific hake) and the metabarcoding data provide information about species composition within a sample (i.e. proportions that sum to one). In combination, these two data sources allow for estimation of DNA concentrations for all species observed by metabarcoding.

##### qPCR model

We developed a state-space framework for modeling hake DNA concentration in the coastal ocean building and modifying the work of Shelton et al. (2022). State-space models separate the true biological process from the methods used to observe the process. Let $D_H(x, y, d)$ be the true, but unobserved concentration of hake DNA (log DNA copies $\mu L^{-1}$) present at spatial coordinates $\{x, y\}$ (eastings and northings, respectively, in km) and sample depth $d$ (meters). Hereafter we refer to a combination of spatial location and depth as a "station" and index station using $j$ (thus, $D_{Hj} = D_H(x, y, d)$). Let $\psi$ be the intercept representing the mean hake DNA concentration and $\epsilon_j$ be a random effect that allows for the effect of that station. We write these as additive on the log-scale,

$$D_{Hj} = \psi + \epsilon_j \tag{1}$$

$$\epsilon_j \sim \text{Normal}(0, \tau) \tag{2}$$

This model describes perhaps the simplest possible random effect model and notably does not leverage any information about the spatial arrangement of samples (i.e. the random effect does not include a spatial correlation component). This is intentional. The focus of this model is on the other species DNA concentration (not Pacific hake) and therefore we allow the estimated hake concentrations at a to follow the qPCR observations as much as possible.

Unfortunately, qPCR does not directly measure DNA concentration. Instead qPCR measures the PCR cycle at which a 2$\mu$L aliquot of a extacted DNA was detected to fluoresce and compares that observation against the flourescence pattern of samples of known DNA concentration ("the standard curve"). This provides an estimate of DNA concentration for each unknown sample. Each water sample is analyzed using at least 3 independent qPCR replicates to account for laboratory and machine variability. In addition to qPCR variability among replicates, we also have to account for modifications that affect each water sample and occurred during water sampling and processing. First, we have three offsets that modify the true DNA concentration to affect what we observed in the qPCR. For water sample $i$: 1) $V_i$ is the proportion of 2.5 L filtered from Niskin $i$ (occasionally some seawater was spilled or not filtered; $V_i = 1$ for the vast majority of samples); 2) $I_i$ is the known dilution used to on sample $i$ to eliminate PCR inhibition. PCR inhibition is most commonly observed in surface samples and is vanishingly rare for samples collected below 100m; 3) $\mathbf{I}\zeta$ is an estimated offset for an ethanol wash error ($\zeta$ is the estimated effect of the wash error and $\mathbf{I}$ is an indicator variable where $\mathbf{I} = 1$ for affected samples and $\mathbf{I} = 0$ otherwise; see Shelton et al. (2022)). Here we fix $\zeta$ = -1.15 based on previous analyses (Shelton et al., 2022).

Finally, we add a random effect for the individual bottles sampled at each station ($\delta_{ij}$). This effect describes the deviation of individual sampled bottles from the location mean, $\delta_{ji} \sim \text{Normal}(0, \sigma_D)$, with $\sigma_D$ indicating a standard deviation among bottles. Because we have only two replicate bottles at each location, we impose a sum to zero constraint on the bottles collected from a single location (i.e. for a given location, $\sum_i \delta_{ji} = 0$). Then, the log-concentration of hake DNA in a sample analyzed by qPCR, $E_{Hji}$ is,

$$E_{Hji} = D_{Hj} + \log V_i + \log I_i + \mathbf{I}\zeta + \delta_{ji} \tag{3}$$

We connect the estimated concentrations to observations from qPCR using two likelihoods connected to the concentration $E$. First we determine whether amplification was detected in each qPCR replicate $r$ run on each plate $p$, $Z_{Hjipr}$ and if amplification was observed we model the PCR cycle at which amplification was detected $Y_{Hjipr}$; $Y_{Hjipr}$ is observed as a continuous, positive value. DNA concentrations have a log-linear relationships with $Y_{Hjipr}$; smaller values of $Y_{Hjipr}$ are associated with higher DNA concentrations.

Because we are modeling the discrete number of DNA molecules that are present in the assayed sample, we know that the Poisson distribution provides an appropriate observation distribution for the number of molecules present in a given qPCR reaction. Assuming a Poisson distribution, conditional on a true mean number of DNA copies in the sample $X$, the probability of having exactly zero DNA copies in a qPCR reaction is $e^{-X}$ and the probability of having non-zero DNA copies is the complement, $1 - e^{-X}$. If there are exactly zero molecules in a qPCR reaction, amplification and therefore detection of amplification will not occur. However, there are other factors (e.g. PCR inhibition) that may reduce amplification further and therefore we expect the probability of amplification to be at most $1 - e^{-X}$. As a result, we estimate an additional term, $\phi$, representing the fractional reduction in amplification efficiency due to other factors during qPCR ($0 < \phi < 1$). Then we have a pair of observation models for hake DNA, both of which are a function of DNA concentration, $E_{Hji}$,

$$Z_{Hjipr} \sim \text{Bernoulli}\left(1 - \exp(-2\phi e^{E_{Hji}})\right) \tag{4}$$

$$Y_{Hjipr} \sim \text{Normal}\left(\beta_{0p} + \beta_{1p} E_{Hji}, \sigma_{Yji}\right) \qquad \text{if } Z_{Hjipr} = 1 \tag{5}$$

The 2 is present in the first line of the equation because we use 2 $\mu$L of sample in each qPCR reaction and so the expected DNA copies in reaction is $2e^{E_{Hji}}$. We allow $\sigma_{Yji}$ to vary as a log-linear function of DNA concentration to account for the fact that there is decreased variability in qPCR measures of $C_t$ at higher DNA concentrations: $\sigma_{Yji} = exp(\gamma_0 + \gamma_1 E_{Hji})$.

By itself, the above model is unidentifiable because field samples do not provide information about the parameters that define relationship between the number of DNA copies and PCR cycle ($\beta_{0j}$, $\beta_{1j}$, and $\phi$). As is standard with qPCR analyses, we include standards of known concentration to estimate these parameters. Each qPCR plate has replicate samples with a known number of DNA copies. These standards span six orders of magnitude (1 to 100,000 copies $\mu$L$^{-1}$, each of 2$\mu$L) and determine the relationship between copy number and PCR cycle of detection. Let $K_{qp}$ be the known log-copy number for sample $q$ in PCR plate $p$, then,

$$Z_{qpr} \sim \text{Bernoulli}\left(1 - exp(-2\phi e^{K_{qp}})\right) \tag{6}$$

$$Y_{qpr} \sim \text{Normal}\left(\beta_{0j} + \beta_{1j} K_{qp}, \sigma_{Yqp}\right) \quad \text{if } Z_{qjp} = 1 \tag{7}$$

Note that there are different intercept ($\beta_{0p}$) and slope ($\beta_{1p}$) parameters for each PCR plate to allow for among-plate variation in amplification. We model each calibration parameter ($\beta_{0p}, \beta_{1p}$) hierarchically using a normal distribution, with among plate mean and variance (i.e. $\beta_{0p} \sim \text{Normal}(\mu_{\beta_0}, \sigma_{\beta_0})$).

##### Metabarcoding model

In addition to the qPCR observations, we used the MiFish primer and high throughput sequencing to detect and enumerate the sequences of other fish species in the same samples that were examined for hake DNA. Again, we are interested in determining the DNA concentration of each focal species in each sample. Unfortunately, metabarcoding data is compositional data (Shelton et al., 2023), and so while it can inform the relative abundance of species within a sample, alone it cannot provide estimates of DNA concentration. Fortunately, the qPCR data from Pacific hake provides information about DNA concentration for one species and the metabarcoding data provides information about the relative abundance for tens of species, including hake. Here, we follow and extend the metabarcoding model developed by Shelton et al. (2023) and show how it links with the qPCR model above.

As above, we let $D_{sj}$ be the log-scale eDNA concentration but now for species $s$. We let $E_{sji}$ be the log-DNA concentration for each species in each bottle. Importantly one of the $s$ species is Pacific hake and so connects to the qPCR model above. Metabarcoding reads are compositional and so we model the DNA concentration relative to one another using additive log-ratio transformation (ARL; Aitchison, 1986). We define a reference species in each sample, $R_i$, with the species used as a reference varying among sample. Let

$\alpha_s$ be the per PCR cycle (43 total number of PCR cycles) log amplification rate for species $s$ (see Shelton et al. (2023)). We can then calculate $\nu_{sji}$, the log-ratio between species $s$ and the reference species after accounting for amplification bias (see *Mock Communities* below),

$$\nu_{sji} = (E_{sji} - E_{Rji}) + N_{PCR}\left(\alpha_s - \alpha_{R_i}\right) \tag{8}$$

We can then convert the $\nu$s to the predicted proportions of the metabarcoding reads $\pi_{sji}$ using the softmax transformation and to the observed reads via a multinomial likelihood,

$$\pi_{sji} = \frac{e^{\nu_{sji}}}{\sum_{s=1}^{S} e^{\nu_{sji}}} \tag{9}$$

$$\boldsymbol{W}_{ji} \sim \text{Multinomial}\left(\boldsymbol{\pi}_{ji}, T_i\right) \tag{10}$$

where $T_i$ is the total number of reads observed across all species for that bottle.

As with the qPCR analyses, in the absence of additional information estimates of $\alpha$ and $E$ cannot be statistically identified. We use the mock community data, which has known input DNA concentrations and same basic model as for the metabarcoding observations but instead of having unknown DNA concentrations, we have known inputs for each species $L_{sm}$ in mock community $m$.

$$\nu_{sm} = (L_{sm} - L_{Rm}) + N_{PCR}\left(\alpha_s - \alpha_{R_i}\right) \tag{11}$$

$$\pi_{sm} = \frac{e^{\nu_{sm}}}{\sum_{s=1}^{S} e^{\nu_{sm}}} \tag{12}$$

$$\boldsymbol{W}_m \sim \text{Multinomial}\left(\boldsymbol{\pi}_m, T_m\right) \tag{13}$$

To maintain identifiability, we assign one $\alpha_s = 0$.

##### Smoothing predictions

The above statistical model provides predictions of species-specific DNA concentrations ($E$) at each depth sampled (d = 0, 50, 150, 300 or 500 m). To generate smooth distribution models, we use posterior means of $E_s(x, y, d)$ from the joint statistical model, exponentiated it to move to concentration from log concentration, $G_s(x, y, d) = exp(E_s(x, y, d))$, and used the sdmTMB package (Anderson et al., 2022) to generate distribution models for the concentration of eDNA. Because of the predominance of very low estimated DNA concentrations at a large number of site, we set $G_s(x, y, d) < 0.5$ equal to zero (SI Appendix Figure S.2) and then used a Tweedie likelihood in sdmTMB,

$$log(F_s(x, y, d)) = \kappa + \eta(x, y) + \omega_d(x, y, d) \tag{14}$$

$$G_s(x, y, d) \sim \text{Tweedie}(F_s(x, y, d), p, \theta_s) \tag{15}$$

where $x$, $y$, and $d$ index longitude, latitude, and depth, respectively. $\kappa$ is scalar intercept. Both $\eta(x, y)$ and $\omega_d(x, y, d)$ are species- and location-specific realizations of Gaussian Markov random fields, $\eta \sim \text{MVN}(0, \sum_\eta)$ and $\omega_d \sim \text{MVN}(0, \sum_{\omega_d})$ (see Anderson et al. (2022)) where $\eta$ is a depth-invariant spatial field and $\omega_d$ is a spatial field specific to each depth category. In simple terms, each location-specific estimate of log eDNA concentration is estimated as a sum of the scalar intercept and two spatial fields, one that is common across depths and one that is depth-specific. $\eta$ estimates the smooth, two-dimensional (latitude/longitude) spatial pattern in estimated DNA, while $\omega_d$ modulates that estimate for different sampling depths. The $\omega_d$ fields were allowed to be independent and identically distributed (IID), and therefore there was no forced correlation across depths beyond that provided by the common spatial field $\eta$. Additionally to convert eDNA concentration from from copies/$\mu$L (reaction concentration) to copies/L of water filtered (ecological concentration) we multiply $F_s(x, y, d)$ by 20 which is the conversion value indicating the ratio of filtered volume and the volume of DNA template used in the PCR reaction.

Estimation of large, multi-level spatiotemporal models can strain computational power, in part because of the need in spatial models to invert large covariance matrices between the data. We use the Stochastic Partial Differential Equation (SPDE) model, implemented through the fmesher R package (Lindgren et al., 2024, 10.32614/CRAN.package.fmesher) to approximate the latent spatial process and its variance by using a triangulated mesh (Lindgren et al., 2011). The SPDE approach drastically reduces computation time by using a sparse covariance matrix to approximate Gaussian Markov random fields like $\eta$ and $\omega_d$. Model estimates and the degree of spatial smoothing can be affected by the details of mesh construction, and we therefore produced a range of meshes with different complexities (i.e., varying numbers of triangles, triangle sizes, and therefore vertices) to test for each species.

Each species was fitted individually using the equation above. We tested 18 candidate meshes for each species, increasing in complexity from 33 to 205 vertices, as well as allowing each model to include or exclude the scalar intercept $\kappa$. Each candidate model for each species was fit by maximum marginal likelihood, implemented in the sdmTMB package in R (Anderson et al., 2022). Model selection was carried out by first ensuring proper convergence, then choosing the model for each species that maximized the summed likelihood.

All the results included in this article are based on the predicted smooth mean concentration, $F_s(x, y, d)$.

#### Model convergance

Convergence and reliability of the Bayesian model were assessed through comprehensive diagnostics. All parameters (SI Appendix Table S.2) exhibited Gelman-Rubin convergence statistics ($\hat{R} < 1.05$) and effective sample sizes (ESS) exceeding 200 per parameter, indicating successful convergence and efficient mixing of the six independent chains (SI Appendix Fig. S.6$A$). An exception was the parameter $\psi$ (mean hake), which showed marginally lower mixing $\text{ESS}_\psi = 149.7$) suggesting mild autocorrelation but no critical failure of convergence.

No divergent transitions were detected during sampling, and the maximum tree depth was not exceeded, indicating no issues with divergence or exploration limits (SI Appendix Fig. S.6$B$). The posterior likelihood demonstrated convergence before the sampling phase began, with all chains exhibiting high mixing, confirming robust exploration of the parameter space (Fig. S.6$B$).

Prior sensitivity analyses revealed that posterior estimates differed from priors, demonstrating that the posteriors were appropriately updated based on the observed data rather than being heavily influenced by prior assumptions (SI Appendix Fig. S.7). Additionally, the posterior predictive checks (PPC) demonstrated that the model reliably reproduced the observed data, supporting the validity of parameter estimates (SI Appendix Fig. S.8).

Collectively, these diagnostics confirm the reliability and validity of the Bayesian model used here.

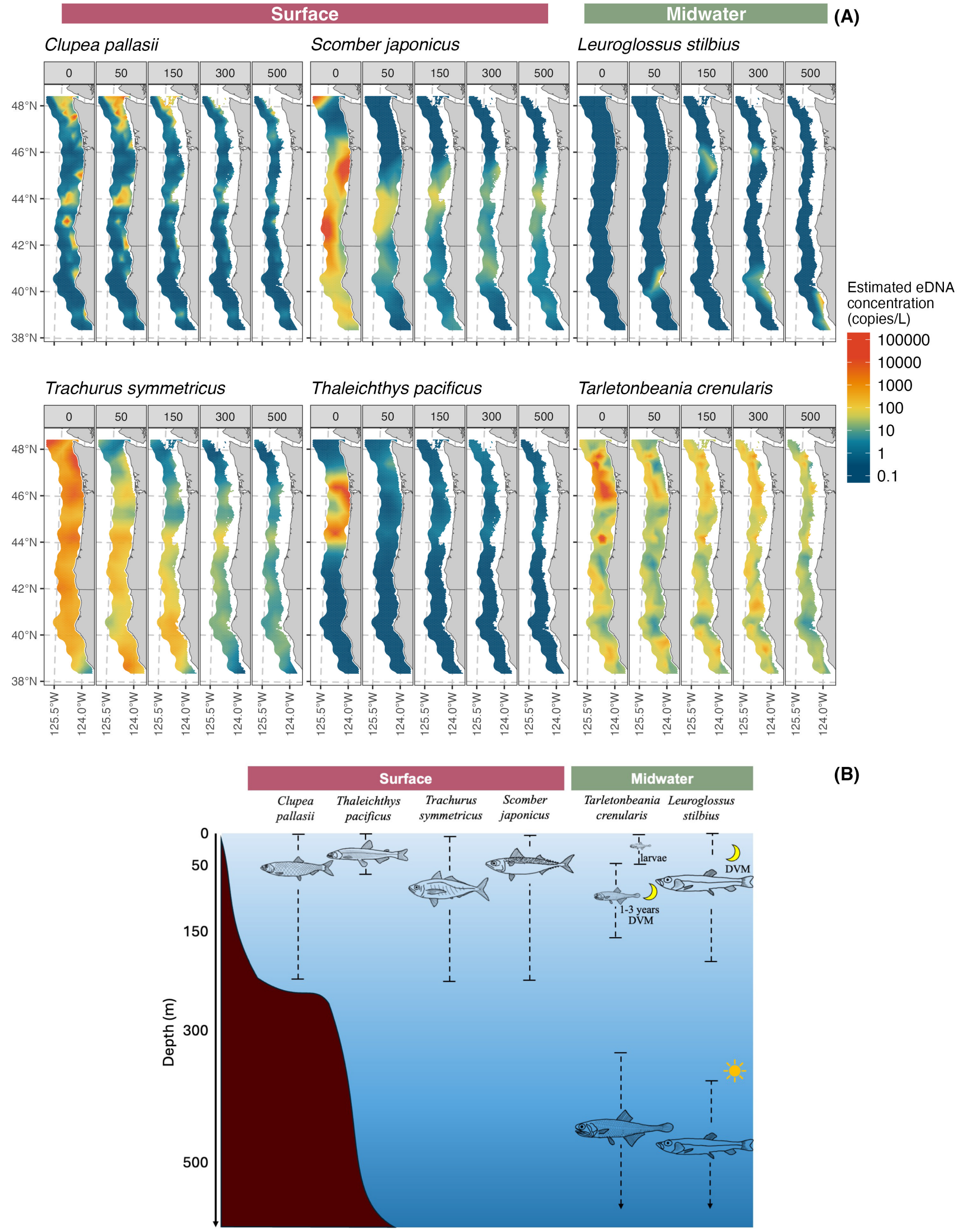


Figure 1: Continuation of Fig. 1. Estimated species eDNA for *Clupea pallasi*, *Trachurus symmetricus*, *Scomber japonicus*, *Thaleichthys pacificus*, *Leuroglossus stilbius*, and *Tarletonbaeania crenularis* concentration (A) across 0, 50, 150, 300, and 500m depth samples and known depth distribution of those species from literature (B).

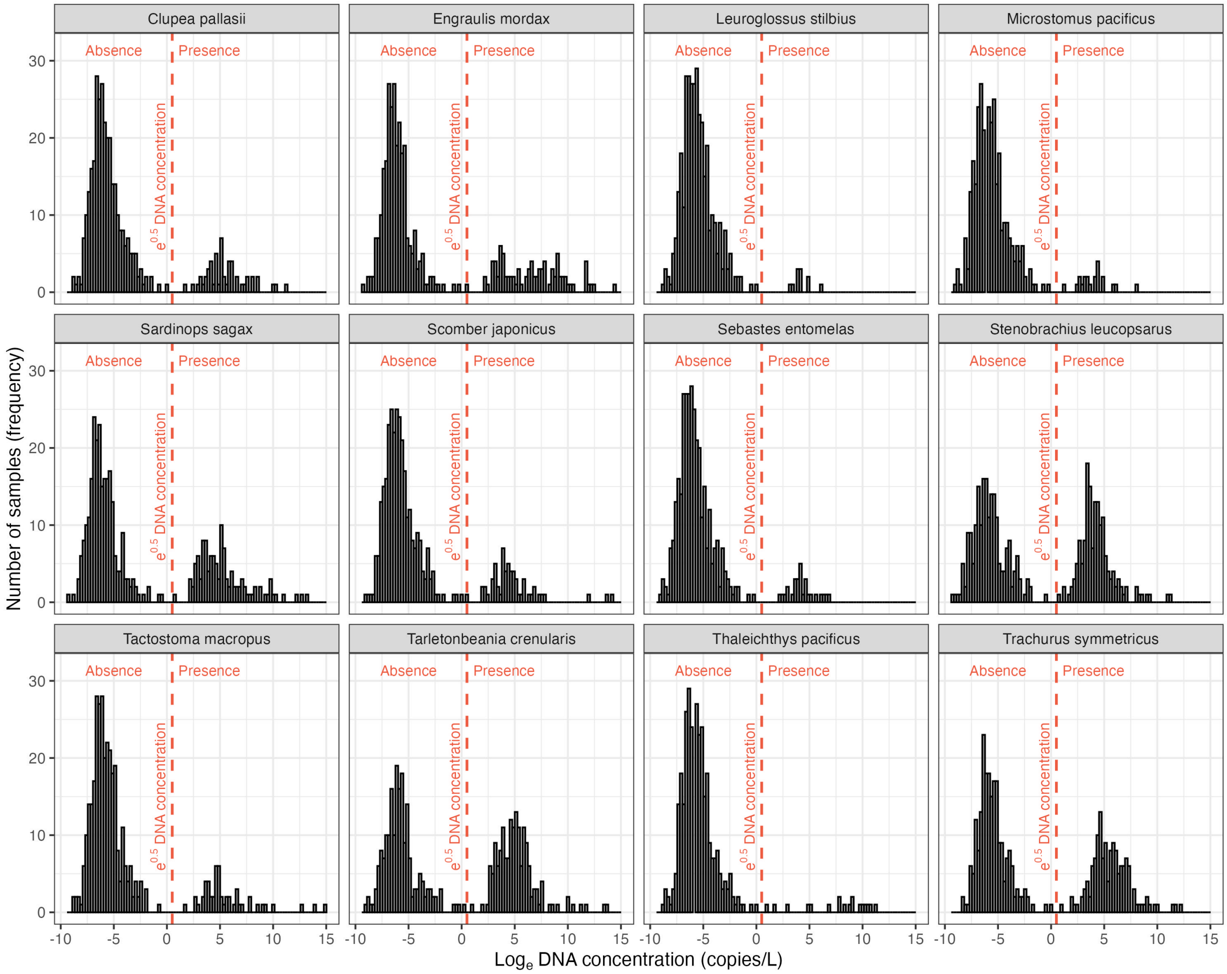


Figure 2: Frequency distribution of eDNA concentration for each species included in this study (n = 12). The dashed red line indicates the threshold model estimated concentration (copies/L) for defining presence and absence of sequenced (through metabarcoding) taxa

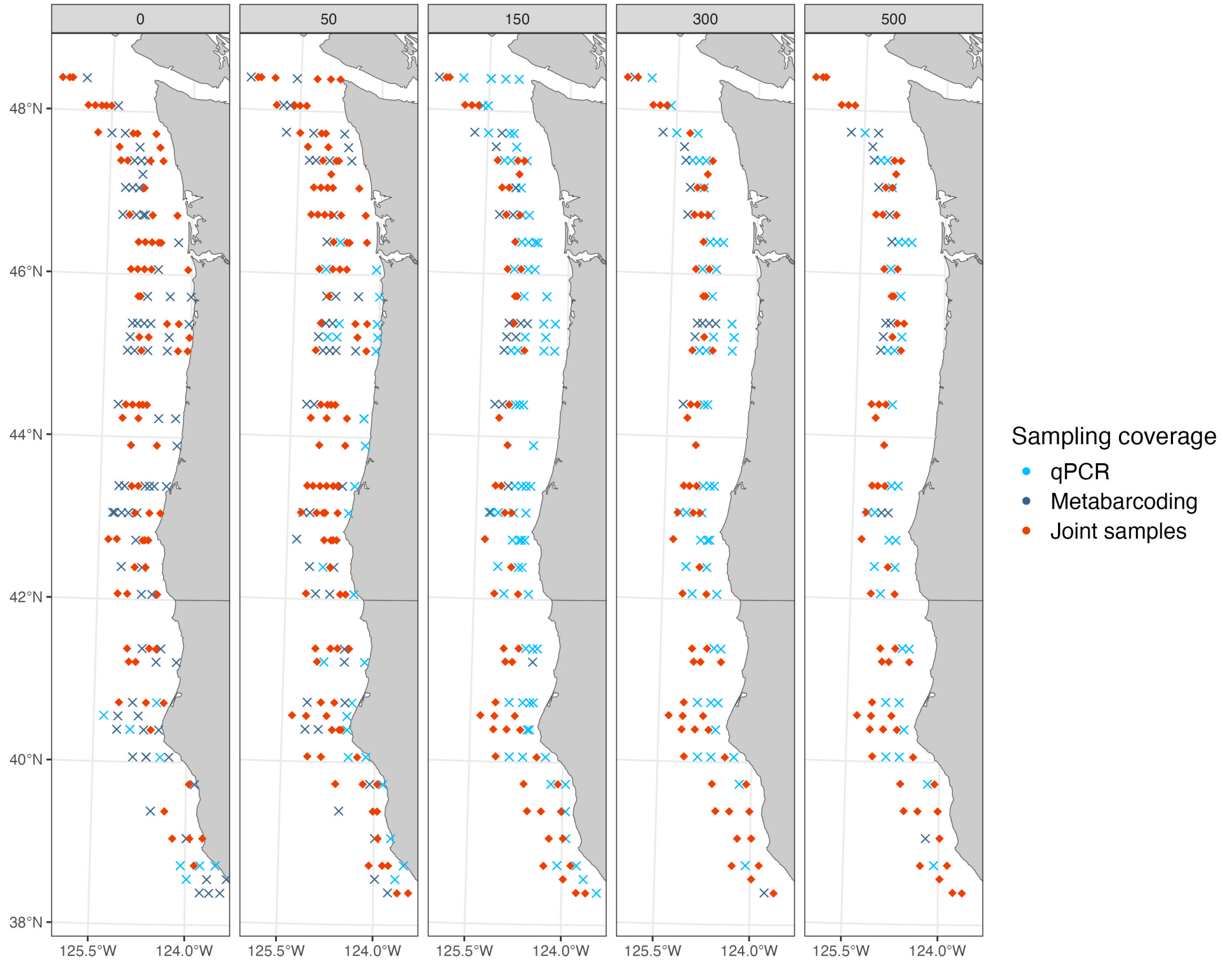


Figure 3: Three-dimensional distribution of samples collected that were run through qPCR (light blue), metabarcoding (dark blue), and were used jointly in the Bayesian model (Fig. 4; red). Jointly modeled samples are those that detected Pacific hake – used as a reference species for quantification – in both qPCR and metabarcoding assays.

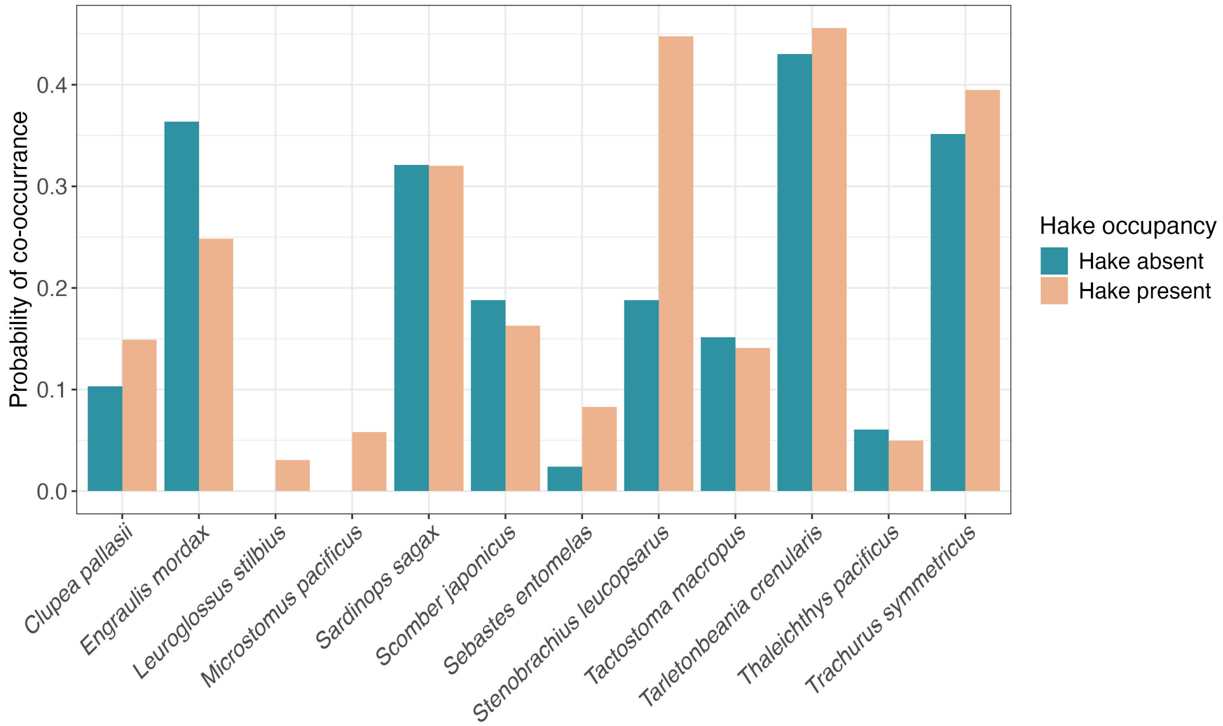


Figure 4: Probability of species co-occurrence with Pacific hake, where blue bars represent non-occurrence and orange bars indicate co-occurrence. *Engraulis mordax* exhibits the most inverse habitat overlap with hake, while *Stenobrachius leucopsarus* shows the highest habitat overlap.

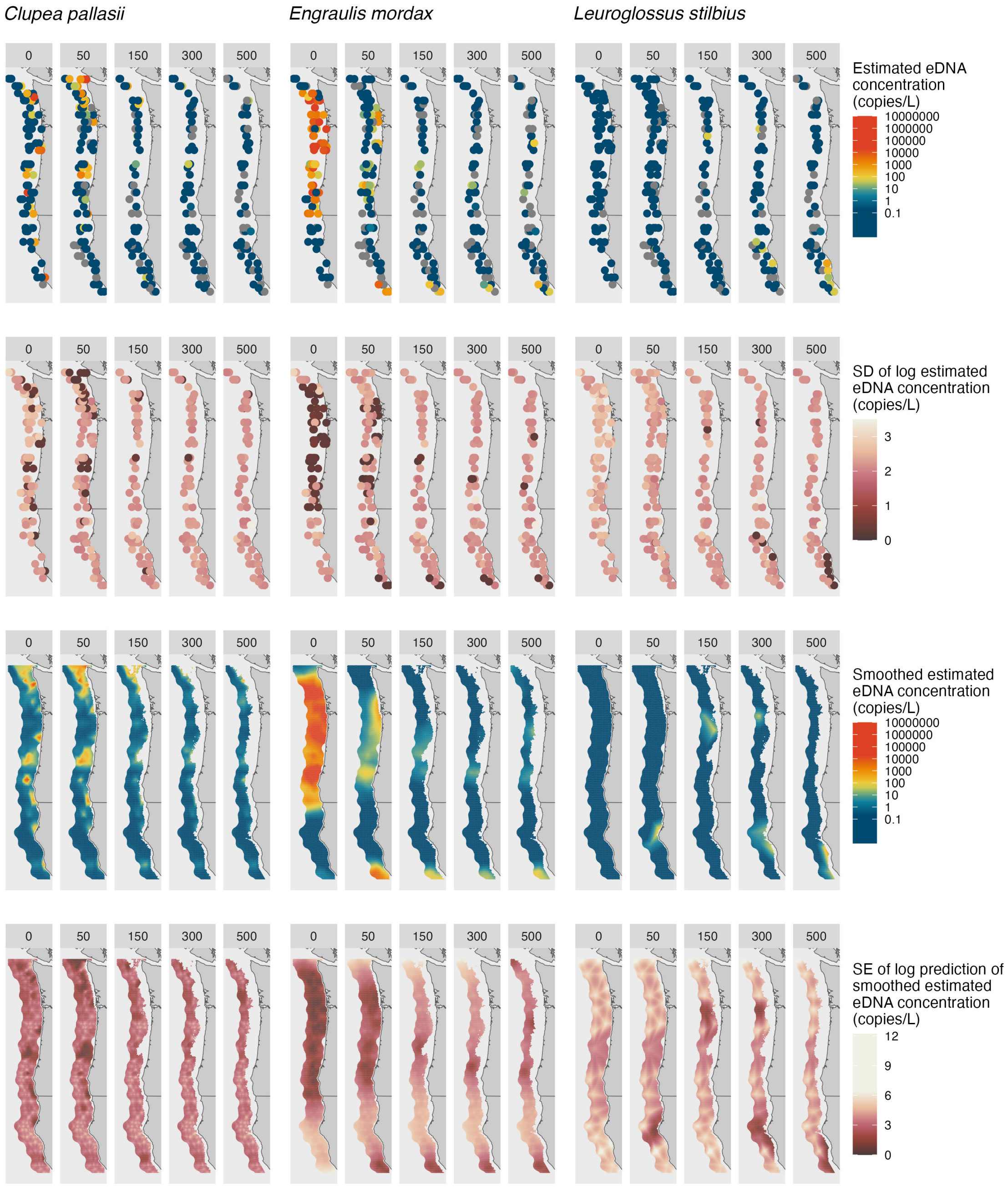


Figure 5: Estimated species (*Clupea pallasii*, *Engraulis mordax*, and *Leuroglossus stiibius*) eDNA concentration (copies/L; first row) through the joint model and their respective standard distribution (SD) of the confidence intervals (second row) across 0, 50, 150, 300, and 500m depth samples. Spatially smoothed eDNA concentration for the same species and depths (third row) with their respective standard error (SE) of prediction.

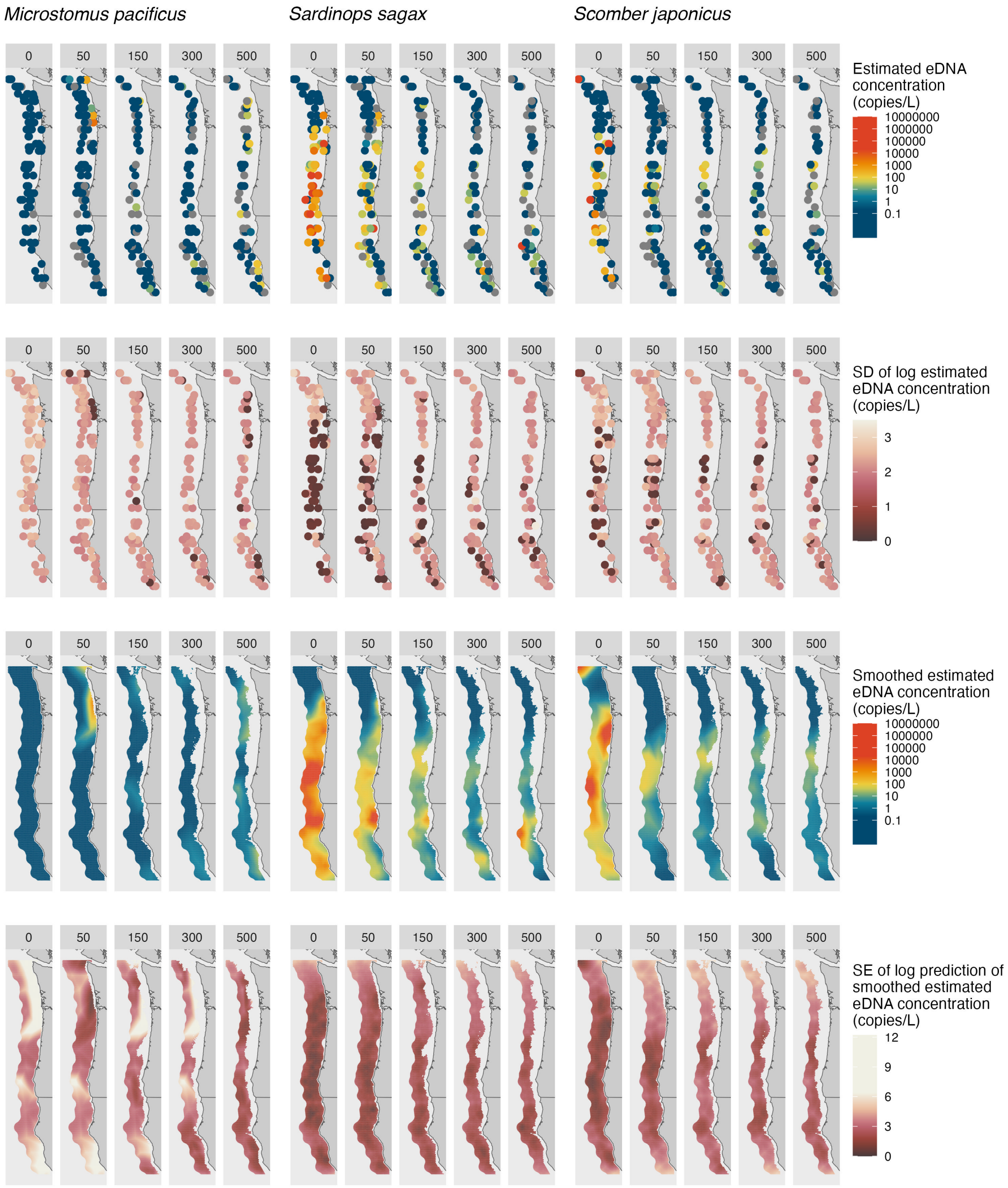


Figure 5 (continued): Species *Microstomus pacificus*, *Sardinops sagax*, and *Scomber japonicus*.

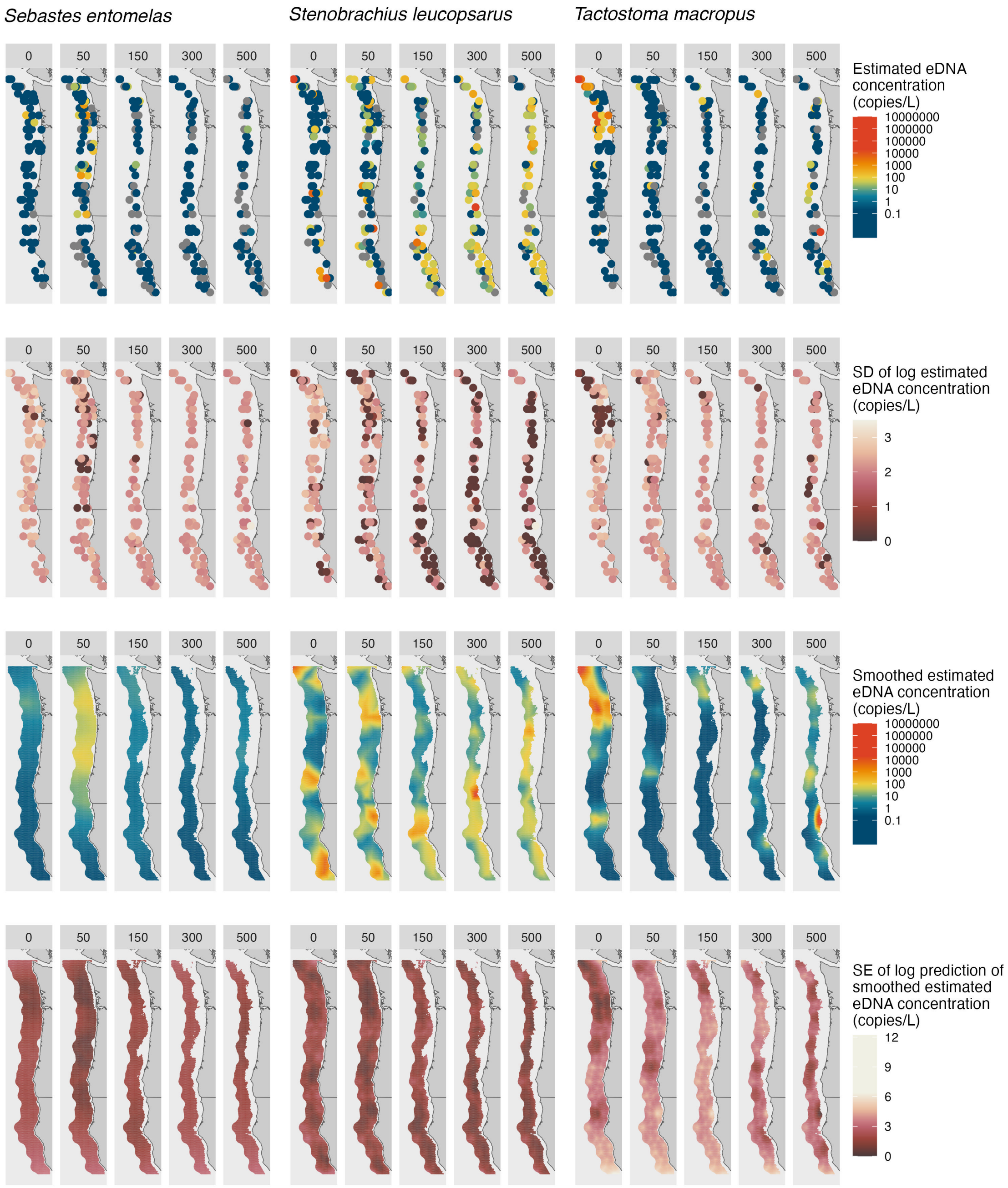


Figure 5 (continued): Species *Sebastes entomelas*, *Stenobrachius leucopsarus*, and *Tacostoma macropus*.

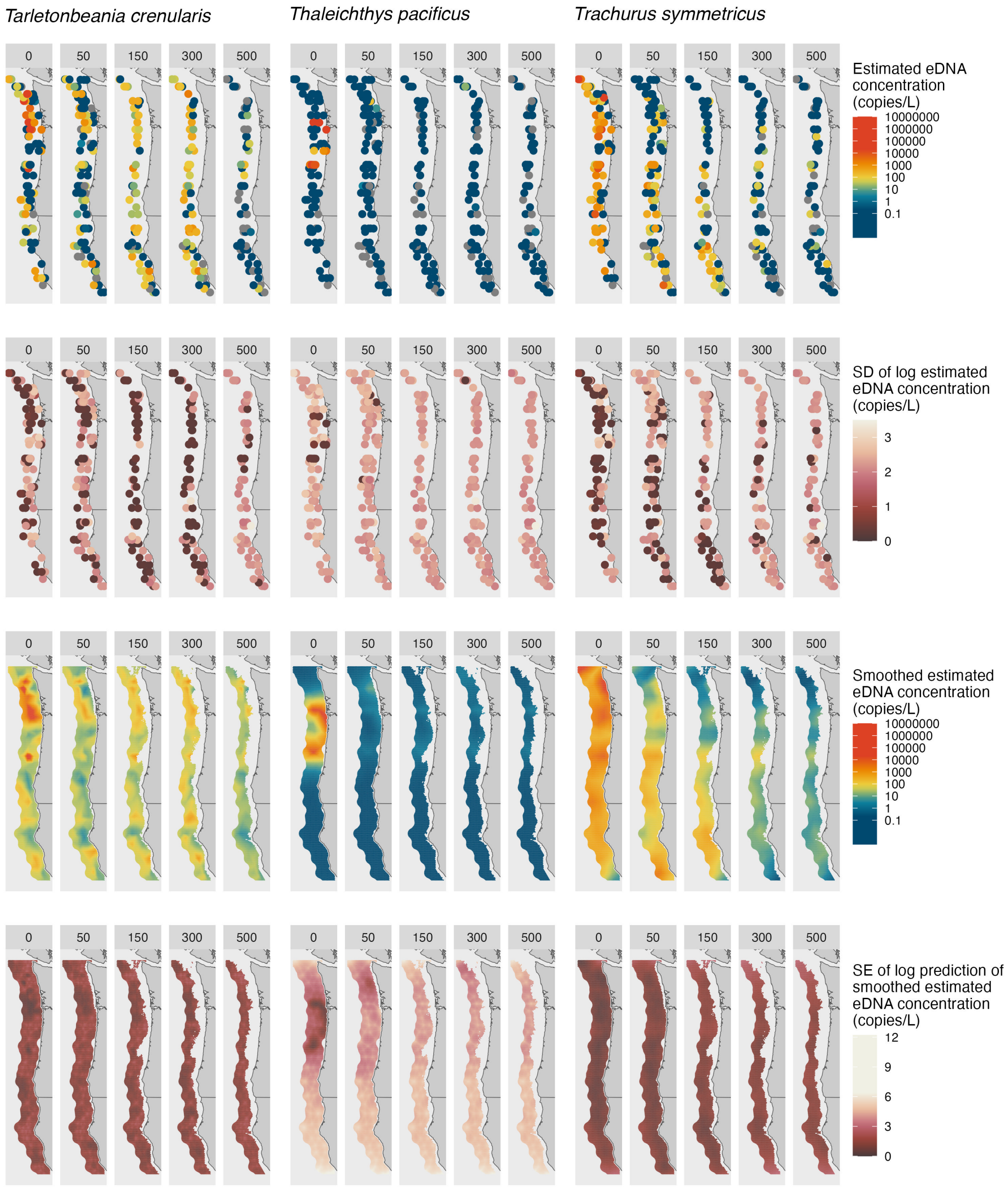


Figure 5 (continued): Species *Tarletonbeania crenularis*, *Thaleichthys pacificus*, and *Trachurus symmetricus*.

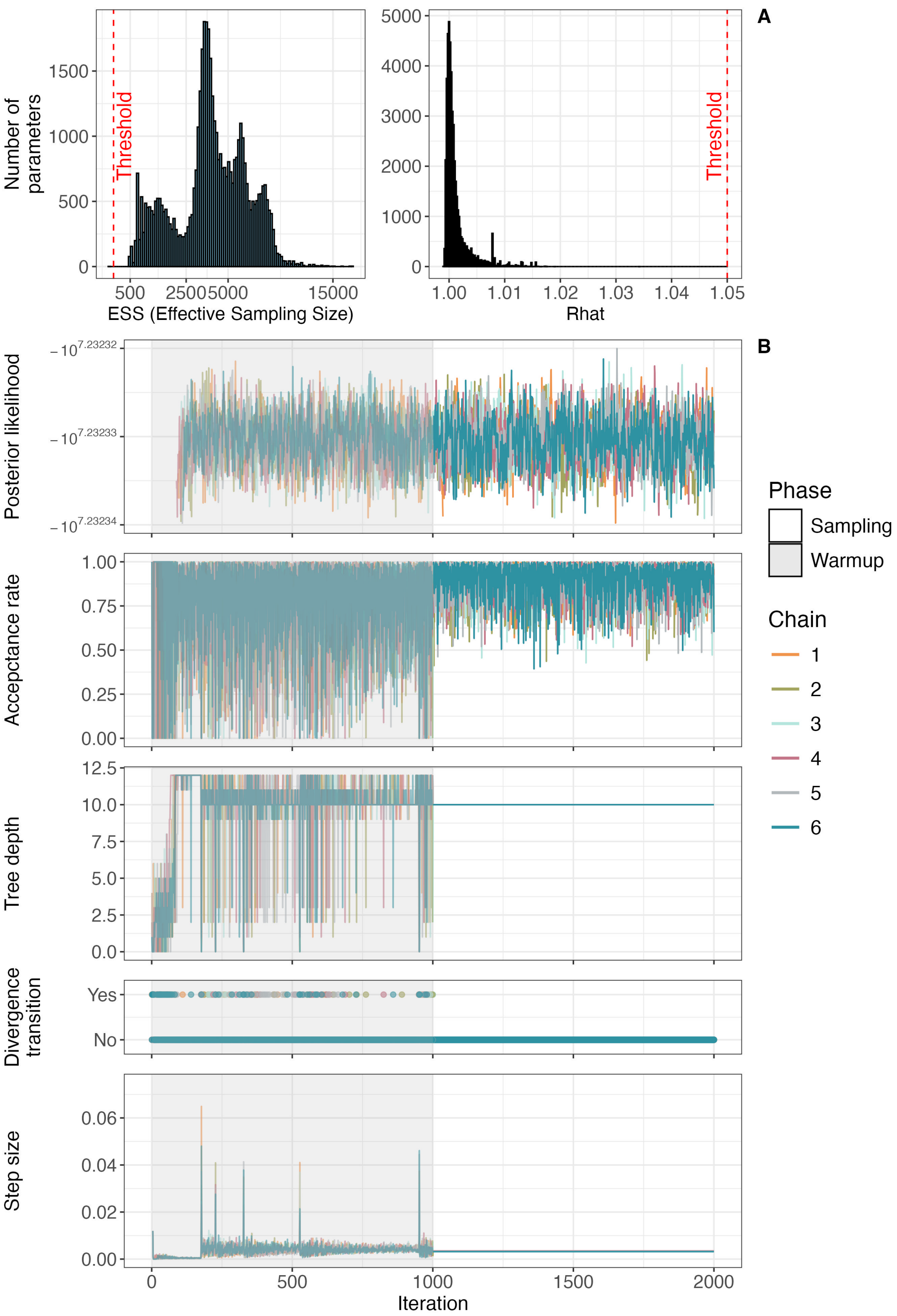


Figure 6: Bayesian model diagnostic plots including $\hat{R}$ and ESS for each parameter (A) and posterior likelihood, acceptance rate, tree depth, divergence transition, and step size for each iteration in each chain (including warm up phase; B).

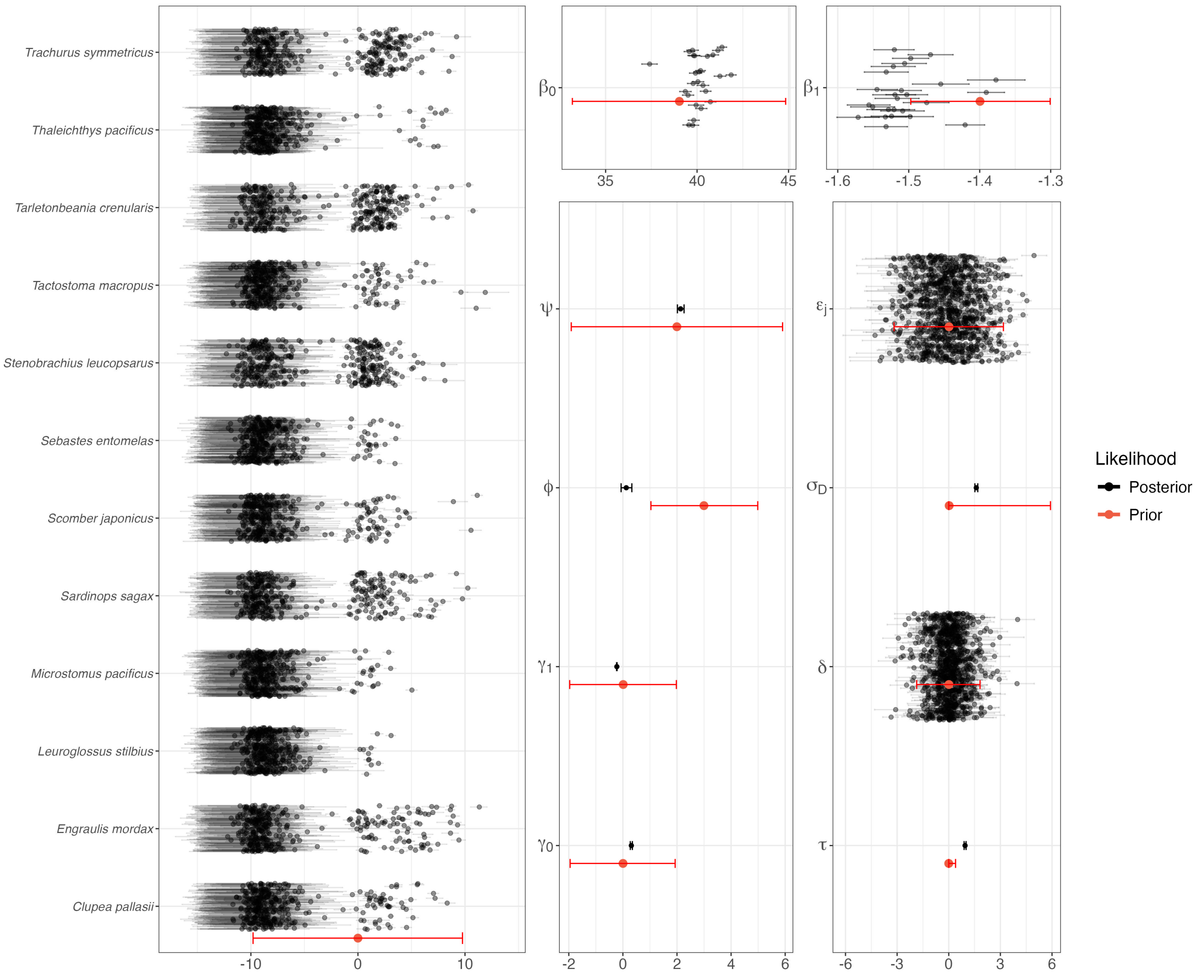


Figure 7: Prior sensitivity plots indicating the distribution of priors in red and posteriors in black with dots indicating the mean and ticks showing 95% credible interval.

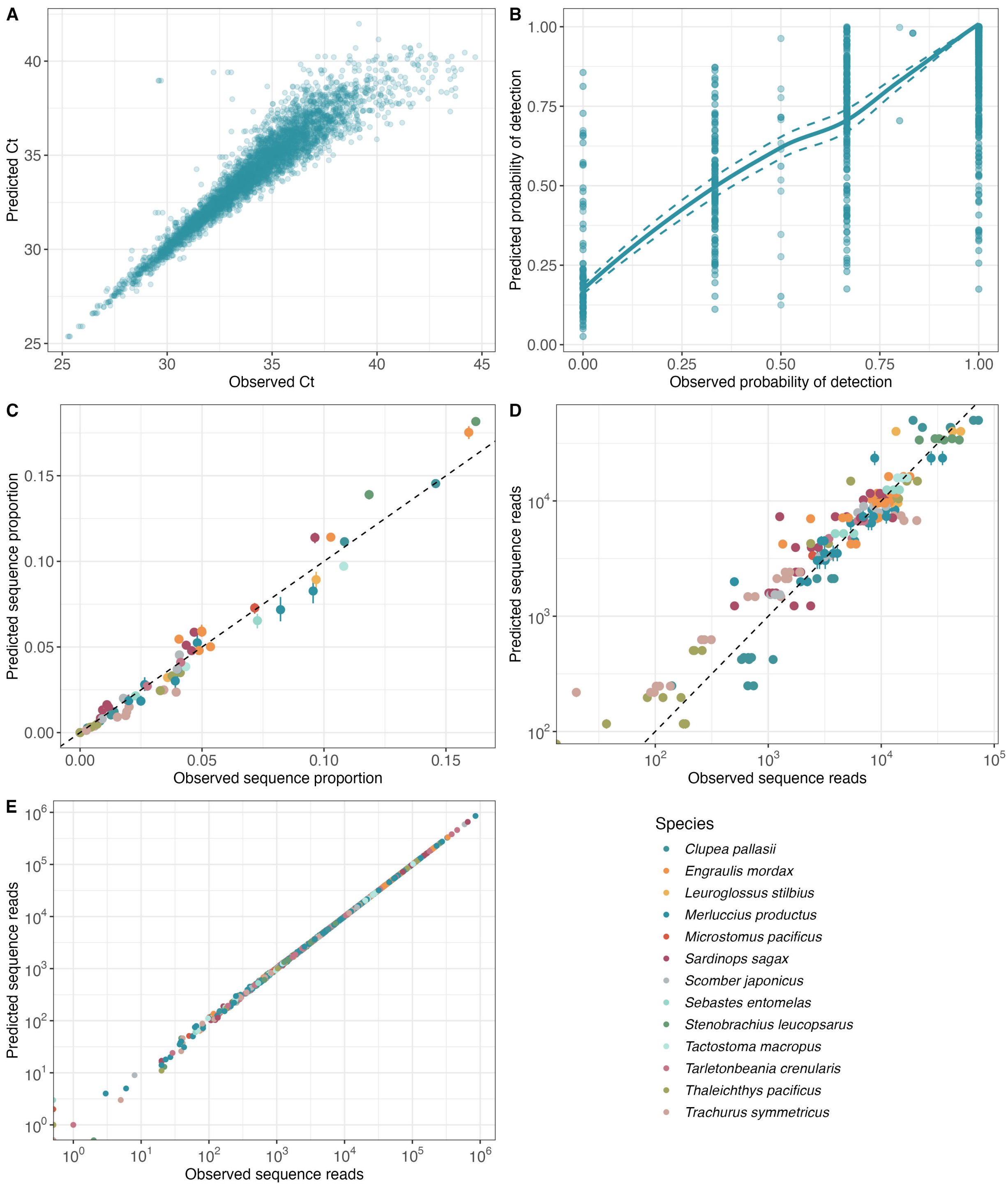


Figure 8: Mean posterior predictions for three model compartments, qPCR continuous model (A), qPCR occupancy model (B), mock community model (C and D on species proportions and number of reads) and metabarcoding model (E)

Table 1: Initial proportional abundances (measured by amplifying the 12S rRNA gene using MarVer1 primers Valsecchi et al. (2020) and ddPCR QX200 Droplet Digital PCR system (Bio-Rad, Inc.)) of the selected species (n=12) across different mock communities (n=8). NA represent no DNA template included.

| Species | Mock1 | | Mock2 | | Mock3 | | Mock4 | |
|---|---|---|---|---|---|---|---|---|
| | Even | Skew | Even | Skew | Even | Skew | Even | Skew |
| *Clupea pallasii* | 0.437 | 0.325 | 0.042 | 0.009 | - | - | 0.025 | 0.038 |
| *Engraulis mordax* | 0.122 | 0.034 | 0.15 | 0.15 | 0.146 | 0.16 | 0.308 | 0.478 |
| *Leuroglossus stilbius* | 0.108 | 0.29 | - | - | - | - | - | - |
| *Merluccius productus* | 0.079 | 0.144 | 0.12 | 0.06 | 0.117 | 0.075 | 0.247 | 0.287 |
| *Microstomus pacificus* | - | - | 0.057 | 0.215 | - | - | - | - |
| *Sardinops sagax* | 0.028 | 0.033 | 0.14 | 0.035 | 0.137 | 0.025 | 0.289 | 0.131 |
| *Scomber japonicus* | - | - | 0.122 | 0.053 | 0.119 | 0.027 | - | - |
| *Sebastes entomelas* | - | - | - | - | 0.068 | 0.218 | - | - |
| *Stenobrachius leucopsarus* | - | - | - | - | 0.356 | 0.487 | - | - |
| *Tactostoma macropus* | - | - | 0.13 | 0.324 | - | - | - | - |
| *Tarletonbeania crenularis* | - | - | 0.083 | 0.124 | - | - | - | - |
| *Thaleichthys pacificus* | 0.123 | 0.113 | 0.099 | 0.012 | - | - | 0.013 | 0.021 |
| *Trachurus symmetricus* | 0.103 | 0.06 | 0.057 | 0.018 | 0.056 | 0.008 | 0.118 | 0.04 |

| | Description | Prior |
|---|---|---|
| | Data | |
| $K$ | known DNA concentration from qPCR stsandards in ln(copies/$\mu$L) | - |
| $Z$ | qPCR amplification (yes=1; no=0) | - |
| $Y$ | qPCR cycle threshold (Ct) | - |
| $R$ | metabarcoding number of reads | - |
| $T$ | total number of reads for all S species | - |
| $N_{PCR}$ | number of metabarcoding PCR cycles | - |
| $L$ | known DNA concentration from mock communinties in ln(copies/$\mu$L) | - |
| $V$ | volume filtered and aliquote dilution offset | - |
| $I$ | aliquote dilution factor | - |
| | State processes | |
| $E_{s\neq H}$ | unknown DNA concentration in ln(copies/$\mu$L) | $\mathcal{N}(0, 5)$ |
| $D$ | unknown DNA concentration in ln(copies/$\mu$L) | - |
| $F$ | smoothed DNA concentration in copies/L | - |
| | Parameters | |
| $\psi$ | overall mean hake eDNA concentration in ln(copies/$\mu$L) | $\mathcal{N}(2, 2)$ |
| $\epsilon_j$ | spatial and depth specific random effect | $\mathcal{N}(0, \tau)$ |
| $\tau$ | standard deviation of biological replicate random effect | $\mathcal{H} - \mathcal{N}(0, 0.2; 0, +\infty)$ |
| $\delta$ | biological replicate random effect | $\mathcal{N}(0, \sigma_D)$ |
| $\sigma_D$ | standard deviation of spatial and depth specific random effect | $\mathcal{H} - \mathcal{N}(0, 3; 0, +\infty)$ |
| $\phi$ | intercept of the qPCR probability of detection relationship | $\mathcal{N}(3, 1)$ |
| $\beta_0$ | intercept of the linear relation between the mean Ct values and eDNA concentration (K and E) | $\mathcal{N}(39, 3)$ |
| $\beta_1$ | slope of the linear relation between the mean Ct values and eDNA concentration (K and E) | $\mathcal{N}(-1.4, 0.05)$ |
| $\gamma_0$ | intercept of the linear relation between the standard deviation of Ct values and eDNA concentration (K and E) | $\mathcal{N}(0, 1)$ |
| $\gamma_1$ | intercept of the linear relation between the standard deviation of Ct values and eDNA concentration (K and E) | $\mathcal{N}(0, 1)$ |
| $\alpha_s$ | species amplification efficiency relative to reference species (R) | fixed (SI Appendix Table S.3) |
| $\zeta$ | ethanol wash effect | fixed (-1.15) |
| | Index | |
| $s$ | species ($H$ = Hake) | - |
| $j$ | location (northing(x), easting(y), depth(d))) | - |
| $i$ | biological replicate | - |
| $r$ | technical replicate | - |
| $p$ | qPCR plate | - |
| $m$ | mock community sample | - |
| $q$ | qPCR standard aliquote sample | - |
| $R$ | metabarcoding reference species | - |

Table 2: Data, state processes, parameters, and subscripts employed in the joint Bayesian model and their prior distributions. $N$ and $H - N$ designate a normal and half-normal distribution, respectively.

Table 3: Species specific metabarcoding amplification efficiency relative to reference species (R; *Merluccius productus* (Pacific hake) here).

| **Species** | $\alpha$ |
|---|---|
| *Clupea pallasii* | -0.00138 |
| *Engraulis mordax* | 0.00557 |
| *Leuroglossus stilbius* | -0.00390 |
| *Microstomus pacificus* | 0.00194 |
| *Sardinops sagax* | 0.00705 |
| *Scomber japonicus* | 0.00434 |
| *Sebastes entomelas* | 0.00459 |
| *Stenobrachius leucopsarus* | 0.00970 |
| *Tactostoma macropus* | -0.00091 |
| *Tarletonbeania crenularis* | 0.00144 |
| *Thaleichthys pacificus* | -0.00506 |
| *Trachurus symmetricus* | -0.00875 |
| *Merluccius productus* | 0.00000 |

### References


Aitchison, J. (1986). *The Statistical Analysis of Compositional Data*. Chapman and Hall, London.

Altschul, S. F., Gish, W., Miller, W., Myers, E. W., and Lipman, D. J. (1990). Basic local alignment search tool. *Journal of Molecular Biology*, 215(3):403–410.

Anderson, S. C., Ward, E. J., English, P. A., Barnett, L. A. K., and Thorson, J. T. (2022). sdmTMB: An R Package for Fast, Flexible, and User-Friendly Generalized Linear Mixed Effects Models with Spatial and Spatiotemporal Random Fields. *bioRxiv*.

Callahan, B. J., McMurdie, P. J., Rosen, M. J., Han, A. W., Johnson, A. J. A., and Holmes, S. P. (2016). DADA2: High-resolution sample inference from Illumina amplicon data. *Nature Methods*, 13(7):581–583.

Guri, G., Shelton, A. O., Kelly, R. P., Yoccoz, N., Johansen, T., Præbel, K., Hanebrekke, T., Ray, J. L., Fall, J., and Westgaard, J.-I. (2024). Predicting trawl catches using environmental DNA. *ICES Journal of Marine Science*, 81(8):1536–1548.

Lindgren, F., Bachl, F., Illian, J., Suen, M. H., Rue, H., and Seaton, A. E. (2024). inlabru: software for fitting latent gaussian models with non-linear predictors. *bioRxiv*.

Lindgren, F., Rue, H., and Lindström, J. (2011). An explicit link between gaussian fields and gaussian markov random fields: The stochastic partial differential equation approach. *Journal of the Royal Statistical Society Series B: Statistical Methodology*, 73(4):423–498.

Martin, M. (2011). Cutadapt removes adapter sequences from high-throughput sequencing reads. *EMBnet.journal*, 17(1):10.

Miya, M., Sato, Y., Fukunaga, T., Sado, T., Poulsen, J. Y., Sato, K., Minamoto, T., Yamamoto, S., Yamanaka, H., Araki, H., Kondoh, M., and Iwasaki, W. (2015). MiFish, a set of universal PCR primers for metabarcoding environmental DNA from fishes: Detection of more than 230 subtropical marine species. *Royal Society Open Science*, 2:150088.

Ramón-Laca, A., Wells, A., and Park, L. (2021). A workflow for the relative quantification of multiple fish species from oceanic water samples using environmental DNA (eDNA) to support large-scale fishery surveys. *PLOS ONE*, 16(9):e0257773.

Shelton, A. O., Gold, Z. J., Jensen, A. J., D'Agnese, E., Andruszkiewicz Allan, E., Van Cise, A., Gallego, R., Ramón-Laca, A., Garber-Yonts, M., Parsons, K., and Kelly, R. P. (2023). Toward quantitative metabarcoding. *Ecology*, 104(2):e3906.

Shelton, A. O., Ramón-Laca, A., Wells, A., Clemons, J., Chu, D., Feist, B. E., Kelly, R. P., Parker-Stetter, S. L., Thomas, R., Nichols, K. M., and Park, L. (2022). Environmental DNA provides quantitative estimates of Pacific hake abundance and distribution in the open ocean. *Proceedings of the Royal Society B: Biological Sciences*, 289(1971):20212613.

Shen, W. and Ren, H. (2021). TaxonKit: A practical and efficient NCBI taxonomy toolkit. *Journal of Genetics and Genomics*, 48(9):844–850.

Valsecchi, E., Bylemans, J., Goodman, S. J., Lombardi, R., Carr, I., Castellano, L., Galimberti, A., and Galli, P. (2020). Novel universal primers for metabarcoding environmental DNA surveys of marine mammals and other marine vertebrates. *Environmental DNA*, 2(4):460–476.
